# FABP8/PMP2 is a positive regulator of PNS myelination

**DOI:** 10.64898/2026.06.26.734879

**Authors:** Jiayue Hong, Clémence Roué, Simar Arora, Ethan McFerland, Sophia Elston, Khushi Panchal, Olivia Harris, Mathieu DiPersio, Taylor Humphrey, Brie Beck, Kaitlin Munro, Senbahavalli Ramasamy, Yannick Poitelon, Sophie Belin

**Author notes:** **Corresponding authors:** Sophie Belin and Yannick Poitelon, Albany Medical College, Department of Neuroscience and Experimental Therapeutics, 47 New Scotland Avenue, MC-136, Albany, NY, 12208. These authors contributed equally to this work (Yannick Poitelon, Sophie Belin).

## Abstract

Myelin formation in peripheral nerves is orchestrated by axon-derived signals and the ability of Schwann cells to expand lipid-rich membrane. Axonal Neuregulin-1 type III (NRG1t3) is known to promote thicker myelin and is associated with increased lipid abundance in peripheral nerves. NRG1t3 also strongly upregulates peripheral myelin protein 2 (FABP8/PMP2), a fatty acid binding protein in myelinating Schwann cells. Here, we directly tested whether elevating PMP2 in Schwann cells is sufficient to enhance myelin growth in nerves. We generated transgenic mice with Schwann cell-specific PMP2 overexpression and showed that PMP2 overexpression drove significant myelin thickening in peripheral nerves, without altering myelin ultrastructure, impairing nerve conduction, or causing detectable adverse effects despite sustained expression. Strikingly, this hypermyelination occurred without activating canonical promyelinating signaling pathways. Instead, elevated PMP2 selectively enhanced fatty acid uptake in Schwann cells, identifying PMP2 as a dosage sensitive enhancer of myelin growth. These findings suggest that in the absence of axonal signaling-driven upregulation, enhanced PMP2 expression itself promotes myelin membrane expansion and enhanced fatty acid availability that could favor complex lipid synthesis. These findings reveal new role for PMP2 as a regulator for myelin expansion *in vivo*, highlighting a key link between lipid trafficking and myelin growth.

## Introduction

The peripheral nervous system (PNS) relies on Schwann cells to myelinate axons and to provide essential metabolic support required for long-term axonal integrity. Myelin formation is a highly coordinated process that depends on precise axon-glia signaling as well as the capacity of Schwann cells to sustain extensive membrane synthesis. Among the axon-derived signals that regulate Schwann cell behavior, Neuregulin-1 type III (NRG1t3) plays a central role in controlling Schwann cell proliferation, differentiation, and the extent of myelin growth during development and in adulthood ^1–3^. Canonically, NRG1t3 signals through ErbB2/ErbB3 receptors to activate downstream pathways such as PI3K-AKT and MAPK-ERK, culminating in the activation of the transcriptional regulator EGR2, a master driver of the myelinating program ^4,5^. Genetic and experimental manipulation of NRG1t3 levels has firmly established that increased axonal NRG1t3 expression promotes myelin thickening during development, after injury, inherited hypomyelinating neuropathies, demyelination, and dysmyelination ^6–8^.

More recently, it has become evident that NRG1t3-driven myelin growth is not solely explained by a proportional upregulation of myelin proteins ^6,7,9^. Sustained elevation of NRG1t3 favors lipid enrichment of the myelin sheath, resulting in thicker myelin with altered lipid composition rather than uniform increases in structural myelin proteins ^6,7,9,10^. In this context, peripheral myelin protein 2 (PMP2; also known as FABP8) has emerged as a uniquely highly responsive downstream effector of enhanced axonal NRG1t3 signaling ^6,7,9^. Unlike other myelin proteins, PMP2 expression is robustly and persistently increased following NRG1t3 overexpression, suggesting that PMP2 may play a specialized role in accommodating the metabolic and biosynthetic demands of enhanced myelin growth ^6,7,9,11^.

PMP2 is a small cytosolic β-barrel protein belonging to the fatty acid–binding protein family and is capable of binding long-chain fatty acids within its hydrophobic pocket. In peripheral nerves, PMP2 displays a striking mosaic pattern of expression, being restricted to a subset of myelinating Schwann cells, preferentially those associated with large-caliber motor axons, and showing variability in expression even along individual internodes ^12–14^. PMP2-positive Schwann cells are characterized by thicker myelin compared with neighboring PMP2-negative cells, implicating PMP2 in myelin growth or maintenance ^6,12,13^. Structural studies have shown that PMP2 can interact with lipid bilayers and bridge membranes at defined spacing, leading to the proposal that PMP2 contributes to myelin assembly or stability ^15–17^. However, genetic loss-of-function studies have challenged a purely structural role for PMP2. Mice lacking *Pmp2* exhibit only transient hypomyelination and modest changes in lipid composition during early development, with largely preserved myelin compaction and nerve conduction later in life ^9,14,18^. Similarly, disease-associated PMP2 mutations causing Charcot-Marie-Tooth disease type 1G (CMT1G) are predicted to retain membrane-binding properties despite their pathogenicity, suggesting that disruption of PMP2 function may involve mechanisms beyond simple defects in myelin formation and compaction ^17,19,20^. Instead, accumulating evidence points to a role for PMP2 in lipid handling and the metabolic adaptation of myelinating Schwann cells ^9,11^.

Myelination is an energetically demanding process that requires the synthesis of large quantities of lipid-rich membrane, making lipid uptake, intracellular transport, and metabolic homeostasis essential for efficient myelin production and long-term maintenance ^21,22^. Schwann cells exhibit remarkable metabolic flexibility, relying on both mitochondrial oxidative phosphorylation and glycolytic pathways, and they actively engage in metabolic coupling with axons by supplying energy substrates such as lactate^23^. Perturbations in Schwann cell metabolism can profoundly affect axonal health without necessarily disrupting myelination itself, highlighting a functional uncoupling between energy production and membrane synthesis ^24,25^. In this context, proteins that coordinate lipid uptake, intracellular trafficking, and utilization are likely to be key regulators of myelin growth. Consistent with this idea, recent works have demonstrated that PMP2 enhances fatty acid uptake, mitochondrial ATP production, and myelin thickening during remyelination ^9,11^. Increased PMP2 expression correlates with elevated incorporation of exogenous long chain fatty acids into Schwann cells and with augmented mitochondrial activity, suggesting that PMP2 may facilitate the delivery of lipid substrates to metabolically active subcellular compartments ^9,11^. However, whether the upregulation of PMP2 is sufficient to drive myelin growth *in vivo* independently of upstream axonal signals such as NRG1t3-mediated PMP2 upregulation, remains unknown. Furthermore, it is unclear whether increased PMP2 expression promotes myelination through activation of canonical promyelinating signaling pathways, modulation of lipid biosynthesis programs, or by acting downstream or in parallel to these pathways through the regulation of intracellular lipid trafficking.

In this study, we tested the hypothesis that PMP2 functions as a dosage-sensitive regulator of Schwann cell myelination. To do so, we generated mice with Schwann cell-specific overexpression of one or two copies of PMP2 (*Pmp2*^OE/+^; P0Cre and *Pmp2*^OE/OE^;P0cre, hereafter referred to as *Pmp2*^OE/+^ and *Pmp2*^OE/OE^), and performed morphometric analyses of peripheral nerves, together with assessments of myelin development, motor and sensory function, and lipid metabolism. Using morphometric, functional, biochemical, and metabolic analyses, we show that elevated PMP2 expression is sufficient to promote sustained myelin thickening during development and adulthood without compromising nerve conduction or activating AKT/ERK or EGR2-dependent transcriptional programs. We further demonstrate that PMP2 overexpression enhances fatty acid uptake through FABP-dependent mechanisms and localizes to metabolically relevant subcellular compartments without inducing endoplasmic reticulum stress or lipid-sensitive transcriptional responses. Together, these findings identify PMP2 as a regulator of Schwann cell fatty acid handling and demonstrate that its increased expression is associated with increased myelin thickness, suggesting PMP2 upregulation as a potential strategy to enhance myelination in disease contexts.

## Methods

### Transgenic mice

All experiments involving animals followed experimental protocols approved by the Albany Medical College Institutional Animal Care and Use Committee. *Pmp2*^OE/+^ and *Pmp2*^OE/OE^ mice were generated with Taconic/Cyagen. A conditional Pmp2 knock-in mouse line was generated by CRISPR/Cas9-mediated genome engineering targeting the ROSA26 locus in C57BL/6J mice. A knock-in cassette consisting of a CAG promoter, loxP-flanked PGK-Neo selection marker, and a Kozak-driven mouse *Pmp2* coding sequence fused to tdTomato (via a 6×EAAAK linker), followed by SV40 and rabbit β-globin polyadenylation signals, was cloned in reverse orientation into intron 1 of the ROSA26 locus. Founder mice were identified by PCR-based genotyping and correct integration was confirmed by sequencing. The genotype of *Pmp2*^OE/+^ and *Pmp2*^OE/OE^ mice was determined by PCR analysis with the *Pmp2*^OE^ allele primers OE-F-5’-CTTTATTAGCCAGAAGTCAGATGC-3’, OE-R-5’-CACAACGAGGACTACACCATCGT-3’ and the *Pmp2^+^* allele primers WT-F-5’-CACTTGCTCTCCCAAAGTCGCTC-3’ WT-R-5’-ATACTCCGAGGCGGATCACAA-3’ that amplified 253bp and 453bp nucleotide fragments for overexpressing and wildtype allele, respectively. PCR conditions were 94⁰C for 5 min, 30 cycles of 94⁰C for 15 sec, and 60⁰C for 30 sec, 72⁰C for 40 sec, followed by 5 min extension at 72⁰C, in a standard PCR reaction mix. The expression of the *Pmp2*-tdTomato transcript is conditional on the expression of Cre recombinase. To target specific expression of Pmp2-tdtomato in Schwann cells, we used a mouse model expressing Cre recombinase under the regulation of the *Mpz* promoter ^26^. *Pmp2*^-/-^ (C57BL/6N-Pmp2tm1(KOMP)Vlcg/MbpMmucd) mice were obtained from the MMRRC. The genotype of *Pmp2*^-/-^ mice was determined by PCR analysis as described previously ^9^. Animals were housed in cages of 5 in 12/12-h light/dark cycles. No animals were excluded from the study. Equal numbers of males and females were included in the study. This study was carried out in accordance with the principles of the Basel Declaration and recommendations of ARRIVE guidelines issued by the NC3Rs and approved by the Albany Medical College Institutional Animal Care and Use Committee (no. 20-08002).

### Behavioral analysis

Grip Strength: Longitudinal grip strength test was performed as previously described ^27^. Mice were placed on a grid connected to an electronic scale. They were gently pulled by the tail with a force parallel to the grid, and they were allowed to grip the grid with all four limbs. The grip strength meter automatically recorded the peak full force generated during each trial before the mouse let go of the grid. This procedure was repeated 10 times per day over two consecutive days for each mouse. The mean values of maximal force across the two days were calculated and normalized to body weight, and the values were reported as grip strength in g. Two-Temperature Preference Test: Two juxtaposed rectangular metal plates were set up underneath the chamber. On day 1, both sides of the metal plates were set at 23⁰C. The mouse was allowed to explore and acclimate for 5 minutes in the chamber. The mouse was subsequently recorded for their time spent on each side using ANY-maze software for a total of 10 minutes. The experiment was repeated on day two, with side I of the metal plates set at 23⁰C, while side II was set at 15⁰C, as previously described ^28^. Von Frey Mechanical Sensitivity Test: Each mouse was acclimated in an individual chamber on a 1x1 cm mesh grid for 30 minutes. Thin filaments of different sizes were used to apply 0.07g, 0.14g, 0.4g, 0,6g, 1.0g, and 2.0g of forces to the hind left paws of the tested mice, as previously described^29^. A rapid withdrawal, a flick, or lick of the stimulated hind paw was recorded as a positive response. Each filament was tested 5 times on each mouse, and the percentages of positive responses at each given force were calculated from those 5 trials.

### Electrophysiological analyses

Mutant and control littermates were analyzed as described previously ^30^. Mice were anesthetized with tribromoethanol 0.4 mg/g of body weight and placed under a heating lamp to avoid hypothermia. Motor action potential recordings of sciatic nerves were obtained with subdermal steel monopolar needle electrodes: a pair of stimulating electrodes were inserted subcutaneously near the nerve at the ankle, then at the sciatic notch, and finally at the paraspinal region at the level of the iliac crest to obtain three distinct sites of stimulation, proximal and distal, along the nerve. Electrophysiological studies comprising motor and sensory nerve conduction studies were conducted using a VikingQuest electromyography device. Measures were blindly evaluated during recordings.

### Morphological analysis

Sciatic nerves were dissected and processed, as described previously ^31^. Nerves were fixed in 2% buffered glutaraldehyde and postfixed in 1% osmium tetroxide. After alcohol dehydration, the samples were embedded in Epon. Transverse sections (0.5–1 nm thick) were stained with toluidine blue and examined by light microscopy. For all morphological assessments, at least three animals per genotype were analyzed. For g ratio analysis of sciatic nerves (axon diameter/fiber diameter), semithin section images were acquired with a ×100 objective. G ratios were determined for at least 100 fibers chosen randomly per animal. Axon and fiber diameters were quantified from semithin sections using the ImageJ software (http://imagej.nih.gov/ij). For myelin periodicity, electron microscopic images were produced by the University at Buffalo Pathology and Anatomical Sciences Research Service Center. Data were analyzed using GraphPad Prism 6.01. Images were blindly evaluated during analysis.

### BODIPY uptake

Mouse sciatic nerve preparation and *ex vivo* fatty acid uptake assays, were performed as described previously ^9^. Briefly, sciatic nerves were harvested, the epineurium was removed, and individual fibers were gently teased apart. Teased fibers were equilibrated in MEM supplemented with 10% lipoprotein-deficient fetal bovine serum at 37°C for 30 min, followed by a 1 h incubation in the same medium containing 10 µM BODIPY-labeled fatty acid C16 (Invitrogen, D3821). Fibers were washed with PBS, mounted on TESPA-coated slides, stained with DAPI, and mounted with ProLong Gold Antifade Mountant (Invitrogen). For FABP inhibition studies, BMS309403 (Tocris) was prepared as a 100 mM stock in DMSO. Rat primary Schwann cells were prepared as previously described ^11^. Schwann cells were pre-incubated overnight with 25 µM BMS309403 or vehicle (0.025% DMSO). On the day of the experiment, cells were equilibrated for 30 min in MEM containing 10% lipoprotein-deficient serum and either inhibitor or vehicle, followed by a 1 h incubation in the same medium supplemented with 10 µM BODIPY-C12 (Invitrogen, D3835). Coverslips were washed with PBS, stained with DAPI, and mounted using ProLong Glass Antifade Mountant (Invitrogen). Sciatic nerve teased fibers and primary Schwann cells were imaged using a Zeiss epifluorescence microscope. Image analysis was performed using ImageJ (https://imagej.nih.gov/ij). For teased fibers, mean fluorescence intensity was measured in three randomly selected regions per fiber. For Schwann cells, total BODIPY fluorescence per field was quantified and normalized to total DAPI counts. Image acquisition and analysis were performed in a blind manner.

### Immunohistochemistry

Teased fibers were prepared as previously described ^32^. OCT-embedded sciatic nerves were sliced into 10 μm-thick sections with cryostat and stored in -80⁰C, as previously described ^33^. Nerve sections and teased fibers were permeabilized with -20⁰C 100% methanol, washed in PBS, then incubated in blocking solution for 1 hr RT. Nerves were incubated overnight at 4⁰C with primary Abs diluted in blocking solution. The following primary Abs were used on cross sections: anti-ChAT 1/100 (Sigma, AB144P), anti-eCadherin 1/100 (Invitrogen, 13-1900), anti-Kdel 1/200 (ENZO Life Sciences, ADI-SPA-827), anti-Neurofilament 1/500 (BioLegend, 822701), anti-P0 1/500 (Aveslabs, PZO), anti-PMP2 1/500 (ProteinTech, 12717-1-AP), anti-tdTomato 1/200 (LSBio, LS-C340696). Nerves were washed, incubated with appropriate secondary Ab diluted in blocking solution for 1 hr at room temperature, washed, stained with DAPI 1/10,000 for 5 min room temperature, washed, mounted with Vectashield, then sealed. Images were acquired with a Zeiss epifluorescent microscope. Analysis was done using ImageJ Software (http://imagej.nih.gov/ij). Images were blindly evaluated during analysis.

### Enzyme-Linked Immunosorbent Assay

The epineurium was removed from the sciatic nerves, and the nerves were flash-frozen in liquid nitrogen, pulverized, and homogenized in 50µl of lysis buffer (50 mM Tris, 100 mM KCl, 12 mM, MgCl_2_ and 1% NP-40). The lysates were incubated on ice for 15 minutes and centrifuged at 15,000 x g for 30 minutes at 4⁰C. Protein concentrations in the supernatants were determined by bicinchoninic acid assay protein assay (Thermo Scientific, Waltham, MA) according to the manufacturer’s instructions. PPARγ activity was measured using the PPAR Complete Transcription Factor Assay Kit (Cayman Chemical, #10008878) according to the manufacturer’s instructions.

### Western Blot

Sciatic nerves were flash-frozen in liquid nitrogen, pulverized, and homogenized in lysis buffer (NaCl 50 mM, HEPES 25 mM, 0.3% CHAPS, pH 7.4, supplemented with 1:100 Protease Inhibitor Cocktail (Roche Diagnostic, Florham Park, NJ), as previously described ^34^. Protein lysates were centrifuged at 15,000 x g for 30 min at 4⁰C. Protein concentrations in the supernatants were determined by bicinchoninic acid protein assay (Thermo Scientific, Waltham, MA) according to the manufacturer’s instructions. Equal amounts of homogenates were diluted 3:1 in 4X Laemmli (Tris–HCl 250 mM, pH 6.8, 8% sodium dodecyl sulfate, 8% β-mercaptoethanol, 40% glycerol, 0.02% bromophenol Blue), denatured for 5 min at 100 °C, resolved on an SDS-polyacrylamide gel, and electroblotted onto PVDF membrane. Blots were blocked with 5% bovine serum albumin in 1X phosphate-buffered saline (PBS), 0.05% Tween-20, and incubated overnight with the following appropriate antibodies: anti-CALNEXIN 1/3,000 (Sigma, C4731), anti-GAPDH 1/3,000 (Sigma, G9545), anti-AKT 1/1,000 (Cell Signaling, 9272), anti-p-AKT Ser473 1/1,000 (9271), anti-ERK1/2 1/1,000 (Cell Signaling, 9102), anti-p-ERK1/2 Thr202/Tyr204 1/1,000 (Cell Signaling, 9101),, anti-MBP 1/1000 (Biolegend, 836504 #SMi-94 and 808401 #SMi-99), anti-PMP2 1/500 (ProteinTech, 12717-1-AP), anti-P0 1/5,000 (Aves Labs, PZO), anti-SREBF2 (ProteinTech, 12943-1-AP), anti-tdTomato 1/1,000 (LSBIO, LS-C340696), anti-β-Tubulin 1/1,000 (Thermofisher, MA5-16309). Membranes were then rinsed in 1X PBS and incubated for 1 hr with secondary antibodies. Blots were developed using ECL or ECL plus (GE Healthcare, Chicago, IL). Western blots were quantified using Image J software (http://imagej.nih.gov/ij).

### RNA preparation and real-time quantitative-PCR

The epineurium was removed from sciatic nerves and the nerves were flash-frozen in liquid nitrogen and pulverized. Total RNA was prepared from sciatic nerve or Schwann cells with TRIzol (Roche Diagnostic). One microgram of RNA was reverse transcribed using Superscript III (Invitrogen) and processed as previously described ^35^. Shortly, for each reaction, 5 ng/μl random hexamers were used. Quantitative PCR was performed using 20 ng of cDNA combined with 1X FastStart Universal Probe Master (Roche Diagnostic) or with TaqMan Real-Time PCR Master Mix (Applied Biosystems), according to manufacturer’s instructions. Data were analyzed using the threshold cycle (Ct) and 2(-ΔΔCt) method. For SyBR Green real time quantitative PCR (RTq-PCR), Rps20 was used as endogenous gene of reference and Rps13 and Rpl27 were used to validate the stable expression of Rps20. Primer sequences were designed using Primer3 and are available in Table S1.

### Statistical Analyses

Experiments were not randomized, but data collection and analysis were performed blind to the conditions of the experiments. Data are presented as mean ± standard error of the mean (s.e.m.) for the in vivo experiments and mean ± standard deviation (s.d.) for the in vitro experiments. Power analysis was used with difference and s.d. commonly observed in the field. Two-tailed Student’s t test, One-way ANOVA and Two-way ANOVA with Bonferroni correction, and linear regression was used for statistical analysis of the differences between multiple groups. Values of p value ≤ 0.05 were considered to represent a significant difference.

## Results

### PMP2 overexpression robustly increases PMP2 in Schwann cells

Our previous studies have shown that PMP2 expression is regulated downstream of axonal Neuregulin-1 type III signaling *in vivo* ^6,7^ and that PMP2 enhances fatty acid uptake, mitochondrial ATP production, and myelination *in vitro* ^9,11^. Conversely, loss of PMP2 alters myelin lipid profiles and transiently impairs nerve conduction ^14,18^. Together, these findings raise the possibility that PMP2 acts as a dosage-sensitive facilitator of metabolic support during myelination.

To directly address this question, we generated mice overexpressing PMP2 specifically in Schwann cells (*Pmp2*^OE/+^ and *Pmp2*^OE/OE^). Using CRISPR/Cas9, we inserted a copy of the *Pmp2* gene fused to a fluorescent tdTomato tag in the ROSA26 locus of the mouse genome. The conditional expression of the PMP2-tdTomato fusion protein is driven by the expression of Cre recombinase under the regulation of the *Mpz* promoter ^26^. Quantitative PCR analysis of sciatic nerves at postnatal day 30 (P30) reveals a gene-dosage–dependent increase in *Pmp2* mRNA (+30% in *Pmp2*^OE/+^ and +65% in *Pmp2*^OE/OE^) when compared with control littermates (Fig. 1A). Consistent with transcriptional upregulation, Western blot analyses demonstrate unchanged level of the endogenous expression of PMP2 (Fig.1D) while showing a robust increase in the PMP2-tdTomato fusion protein in *Pmp2*^OE/+^ and *Pmp2*^OE/OE^ nerves at both P30 and adulthood (P240) (Fig. 1B-E).

**Figure 1.**
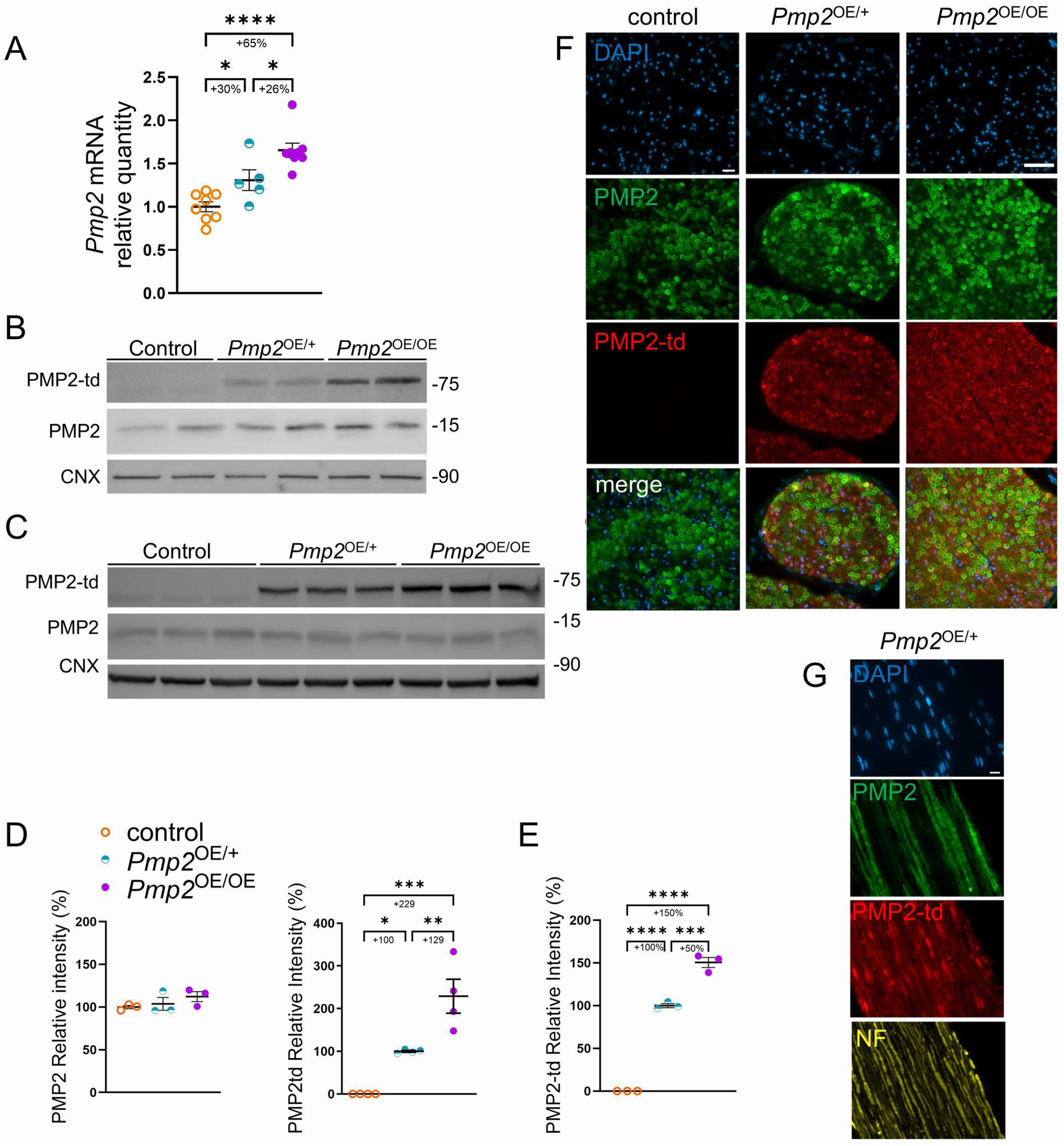
Quantification of PMP2 overexpression in PMP2 overexpressing mouse sciatic nerves. (A) qPCR analysis for *Pmp2* mRNA level in control, *Pmp2*^OE/+^, *Pmp2*^OE/OE^ mouse sciatic nerves at P30. *Rpl27* was used as a housekeeping control. (B-C) Western blot analysis and (D-E) densitometry for PMP2 and PMP2-tdTomato in control, *Pmp2*^OE/+^ and *Pmp2*^OE/OE^ sciatic nerves at P30 (B, D) and P240 (C, E). Calnexin (CNX) was used as a protein loading control. PMP2-td (PMP-tdTomato). (F) Epifluorescent imaging of PMP2 (green), PMP2-tdTomato (red), and DAPI (blue) in control, *Pmp2*^OE/+^, and *Pmp2*^OE/OE^ cross sections from mouse sciatic nerve at P30. (F) Epifluorescent imaging of PMP2 (green), PMP2-tdTomato (red), and Neurofilament (NF, yellow) in *Pmp2*^OE/+^ longitudinal sections from mouse sciatic nerve at P30. Scale bars = 10 µm. Error bars represent s.e.m. n = 3-8 mice, and each data point represents a different n. One-way ANOVA with Bonferroni post hoc test. *p<0.05, **p<0.01, ***p<0.001, ****p<0.0001.

Co-immunostaining for endogenous PMP2 and the PMP2-tdTomato transgenic fusion protein reveals extensive colocalization, with merged images showing yellow-orange overlap in the majority of cells across in *Pmp2*^OE/+^ and *Pmp2*^OE/OE^ sciatic nerve cross sections (Fig. 1F). However, the two signals are not entirely superimposable: endogenous PMP2 exhibited a sharply defined compact myelin staining, whereas PMP2-tdTomato display a more diffuse, punctate cytoplasmic distribution that extended beyond the boundaries delineated by the endogenous protein. Longitudinal sections reveal additional enrichment of the PMP2–tdTomato fusion protein in Schwann cell nuclei (Fig. 1G).

Overall, these data indicate that *Pmp2*^OE/+^ and *Pmp2*^OE/OE^ expressed higher level of PMP2 than control mice.

### Elevated PMP2 modulates Schwann cell–axon interactions

Because PMP2 expression is regulated by axonal cues, and PMP2 levels in Schwann cells are tightly correlated with motor axons ^12,13^, we next asked whether elevated PMP2 alters Schwann cell–axon interactions. Indeed, changes in PMP2 levels in Schwann cells could influence Schwann cells’ myelinating fate, particularly for axonal subtypes with high energetic and lipid demands ^22,36^.

Quantitative analysis of sciatic nerve cross sections reveals a significant increase in the proportion of PMP2-positive Schwann cells in *Pmp2*^OE/+^ and *Pmp2*^OE/OE^ nerves, with an increase of 68% and 44%, respectively, when compared to control nerves (Fig. 2A-B). The proportion of ChAT-positive axons (a marker of motoneurons) myelinated by PMP2-positive Schwann cells increases in *Pmp2*^OE/+^ and *Pmp2*^OE/OE^ nerves by 45% and 37% respectively, when compared to control nerves (Fig. 2C). However, when normalized to the total PMP2-positive population, a lower fraction of PMP2-positive Schwann cells engages ChAT-positive axons in *Pmp2*^OE/+^ compared with controls (Fig. 2D). This suggests that the elevated myelination of motor axons in Fig. 2C reflects an overall increase in PMP2-positive Schwann cells rather than a preferential targeting of cholinergic axons. This lack of specificity is expected, as PMP2 overexpression is driven by P0-Cre, which recombines in all Schwann cells, regardless of the identity of the axon, rather than being induced by an axon-derived cue specific to motor neurons.

**Figure 2.**
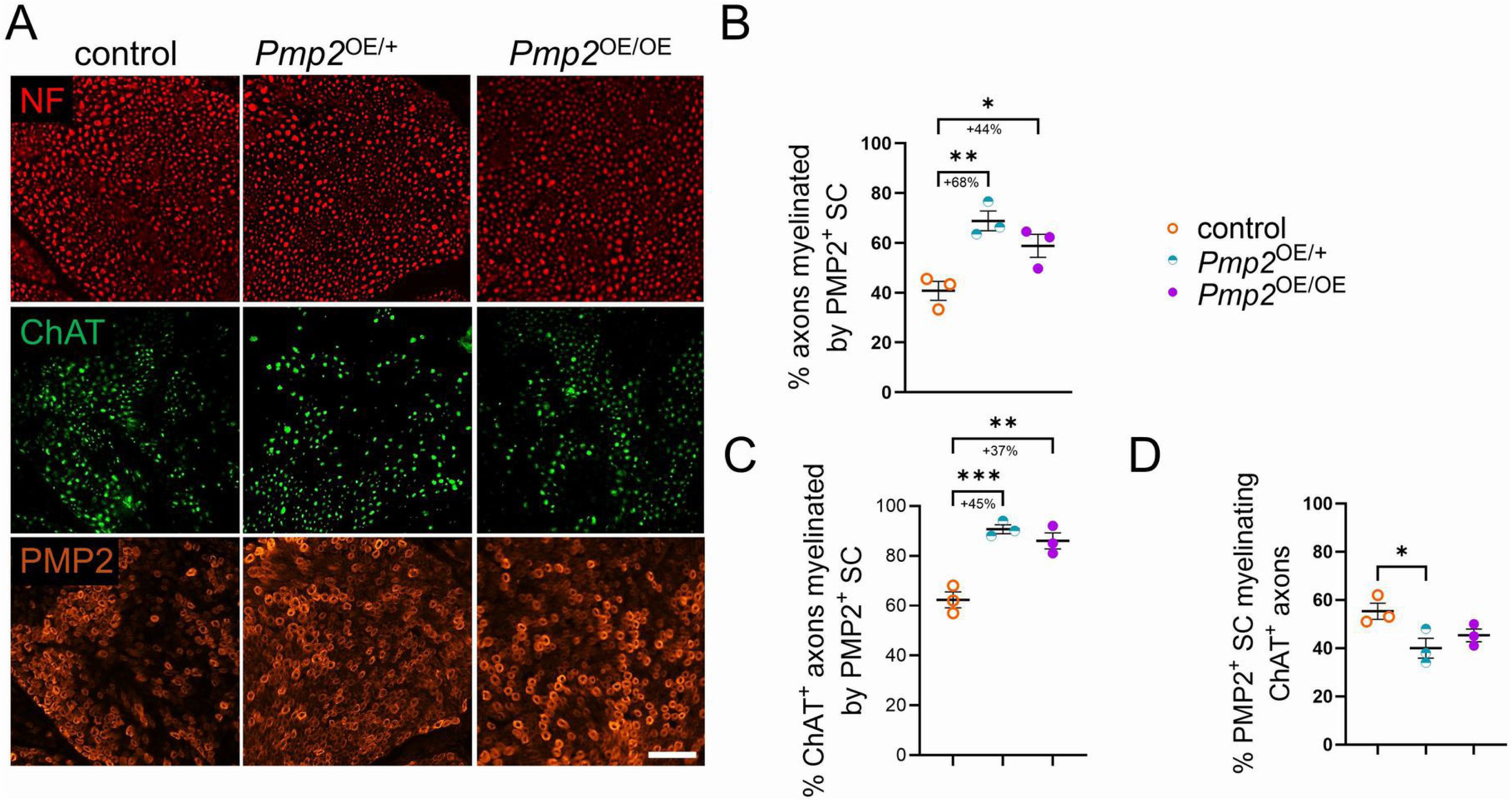
Quantification of PMP2-expressing Schwann cells and ChAT-expressing axons in PMP2 overexpressing mouse sciatic nerves at P30. (A) Epifluorescent imaging of PMP2 (orange), ChAT (green), and Neurofilament (NF, red) in control, *Pmp2*^OE/+^ and *Pmp2*^OE/OE^ cross sections from mouse sciatic nerve at P30. Scale bars = 50 µm. (B) Percentage of PMP2-positive (PMP2^+^) Schwann cells, normalized to NF. (C) Percentage of ChAT^+^ axons myelinated by PMP2^+^ Schwann cells, normalized to ChAT^+^ axons. (D) Percentage of PMP2^+^ Schwann cells myelinating ChAT^+^ axons, normalized to PMP2^+^ Schwann cells. Error bars represent s.e.m. n = 3 mice, and each data point represents a different n. One-way ANOVA with Bonferroni post hoc test. *p<0.05, **p<0.01, ***p<0.001.

Together, these data indicate that elevated PMP2 expands the pool of myelinating Schwann cells and increases their overall axon engagement capacity of both ChAT-positive and non-ChAT-positive axons.

### Sustained PMP2 overexpression in Schwann cells increases myelin thickness in development and adulthood

Because elevated PMP2 expression leads to more PMP2-positive Schwann cells in *Pmp2*^OE/+^ and in *Pmp2*^OE/OE^ mice sciatic nerves, and given PMP2’s role as fatty acid binding protein and its known effects in lipid handling and myelin formation *in vitro* ^9,11^, we next asked whether more PMP2-positive Schwann cells are correlated with an increase in myelin thickness in sciatic nerves.

Morphometric analysis of semithin and ultrastructural sciatic nerve sections at P30 revealed a significant reduction in g ratios in *Pmp2*^OE/+^ a mice sciatic nerves compared with controls (Fig. 3A-B). This effect was most pronounced in small- and medium-caliber axons (<4 μm diameter), whereas large-caliber fibers showed non-significant changes; possibly because large axons have higher baseline PMP2. Notably, this reduction in g ratio was not observed in *Pmp2*^OE/OE^ mice sciatic nerves, suggesting that the relationship between PMP2 dosage and myelin thickening is non-monotonic at this early postnatal timepoint. Importantly, axon diameter distributions were not significantly altered across genotypes, indicating that the reduced g-ratio reflects increased myelin thickness rather than axonal hypotrophy (Fig. 3C). By 8 months, the *Pmp2*^OE/+^ phenotype persisted, and *Pmp2*^OE/OE^ mice now also show significantly reduced g ratios across all axon diameters (Fig. 3D-F). Ultrastructural analysis by electron microscopy done at 6 months demonstrated normal myelin compaction, lamellar organization, and periodicity in *Pmp2*^OE/+^ compared to control sciatic nerves (Fig. 3G, H), indicating that PMP2 overexpression promotes sustained myelin growth without disrupting myelin ultrastructure.

**Figure 3.**
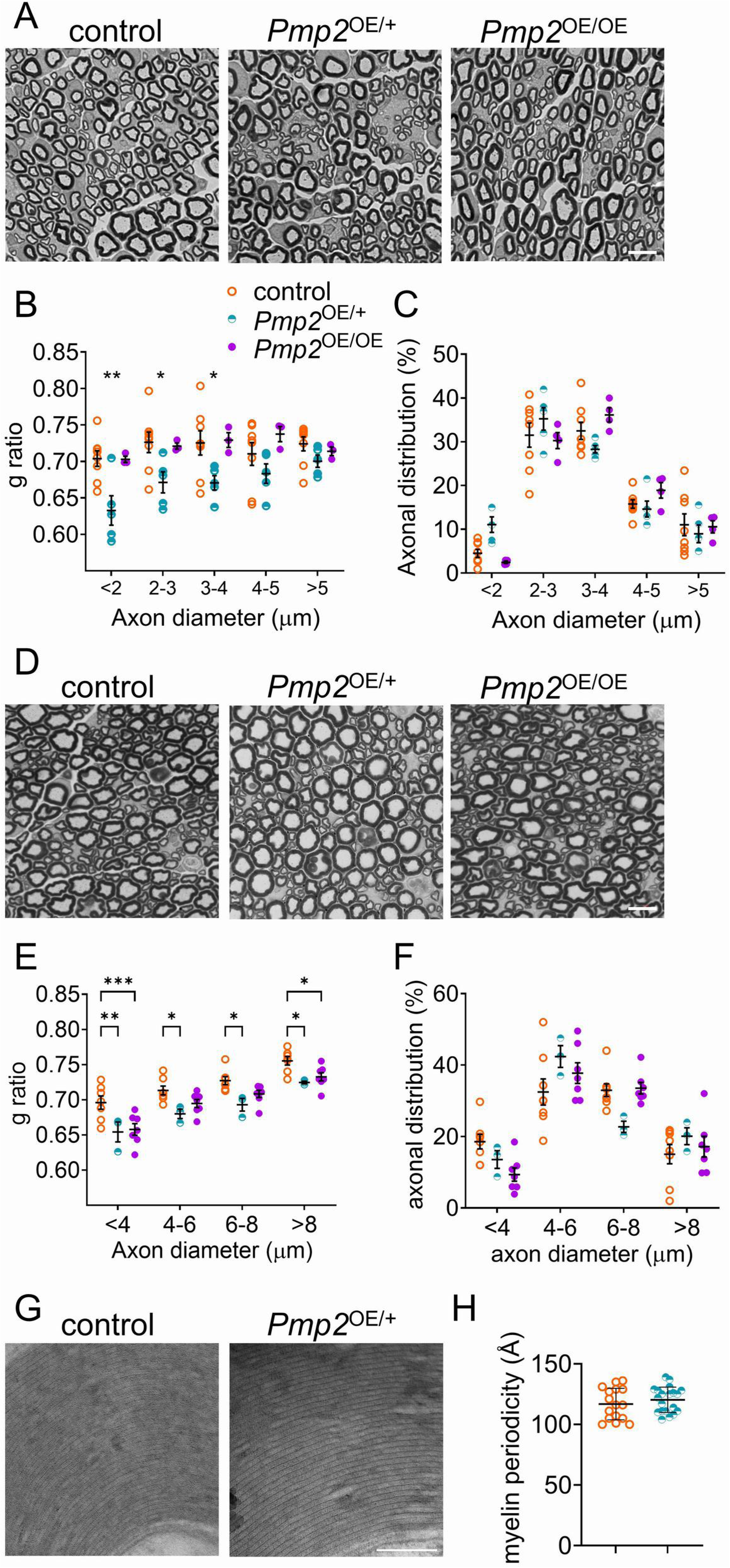
Morphological analysis of PMP2 overexpressing mouse sciatic nerves. (A, D) Semithin sections of control, *Pmp2*^OE/+^, and *Pmp2*^OE/OE^ mouse sciatic nerves at P30 (A) and P240 (D). Scale bar = 10µm. (B, E) Myelin thickness, and (C, F) distribution of myelinated axon diameters. Error bars represents s.e.m. Each point represents a different mouse (B, C, E, F). n = 4-8 mice. Two-way ANOVA with Bonferroni post hoc test. *p<0.05, **p<0.01, ***p<0.001. (G) Electron micrographs of control and *Pmp2*^OE/+^ mouse sciatic nerves at P160 (H) Myelin periodicity. Scale bar = 1µm. Each point represents a different fiber (H). n = 16-20.

Because, increased myelin thickness could be associated with altered conduction properties or neuropathic phenotypes ^37^, we assessed whether the structural changes observed in *Pmp2*^OE/+^ and in *Pmp2*^OE/OE^ nerves were accompanied by functional deficits. Longitudinal grip strength measurements from P30 to P240 revealed no significant differences between genotypes (Supplementary Fig. 1A). Electrophysiological recordings at P240 showed normal compound muscle action potential amplitudes, nerve conduction velocities, and F-wave latencies (Supplementary Fig. 1B). Sensory behaviors, including thermal and mechanical sensitivity, were also preserved (Supplementary Fig. 1C, D). Thus, despite inducing substantial structural remodeling of myelin, PMP2 overexpression does not compromise peripheral nerve function.

Given that increased myelin thickness at P30 was observed in *Pmp2*^OE/+^ but not in *Pmp2*^OE/OE^ nerves, we next asked whether this dosage-dependent dissociation was reflected in canonical promyelinating signaling pathways, examining the expression and activation of key regulators of Schwann cell differentiation and myelination ^38^. Quantitative RT-PCR analysis revealed no significant differences in *Egr2* mRNA levels between control, *Pmp2*^OE/+^, and *Pmp2*^OE/OE^ sciatic nerves (Fig. 4A). Consistent with this, protein levels of the major myelin proteins P0 and MBP were unchanged across genotypes (Fig. 4B), indicating that PMP2 overexpression does not globally enhance the core myelination program. We next assessed activation of AKT and ERK signaling pathways, which are critical mediators of axon-dependent Schwann cell myelination ^38^. Western blot analyses showed no significant differences in total or phosphorylated AKT and ERK levels in *Pmp2*^OE/+^ and *Pmp2*^OE/OE^ nerves compared with controls (Fig. 4C), suggesting that PMP2-driven increases in myelin thickness in *Pmp2*^OE/+^ occur independently of sustained changes in these signaling pathways. Q-PCR analysis further revealed that genes involved in fatty acid and lipid biosynthesis were largely unchanged in *Pmp2*^OE/+^, and *Pmp2*^OE/OE^ sciatic nerves (Supplementary Fig. 2). While *Srebf2* mRNA was modestly increased in *Pmp2*^OE/OE^ nerves but not in *Pmp2*^OE/+^, this did not translate into increased SREBF2 protein abundance, indicating that lipid biosynthetic capacity might not be modulated at the protein level by the overexpression of PMP2 in Schwann cells.

**Figure 4.**
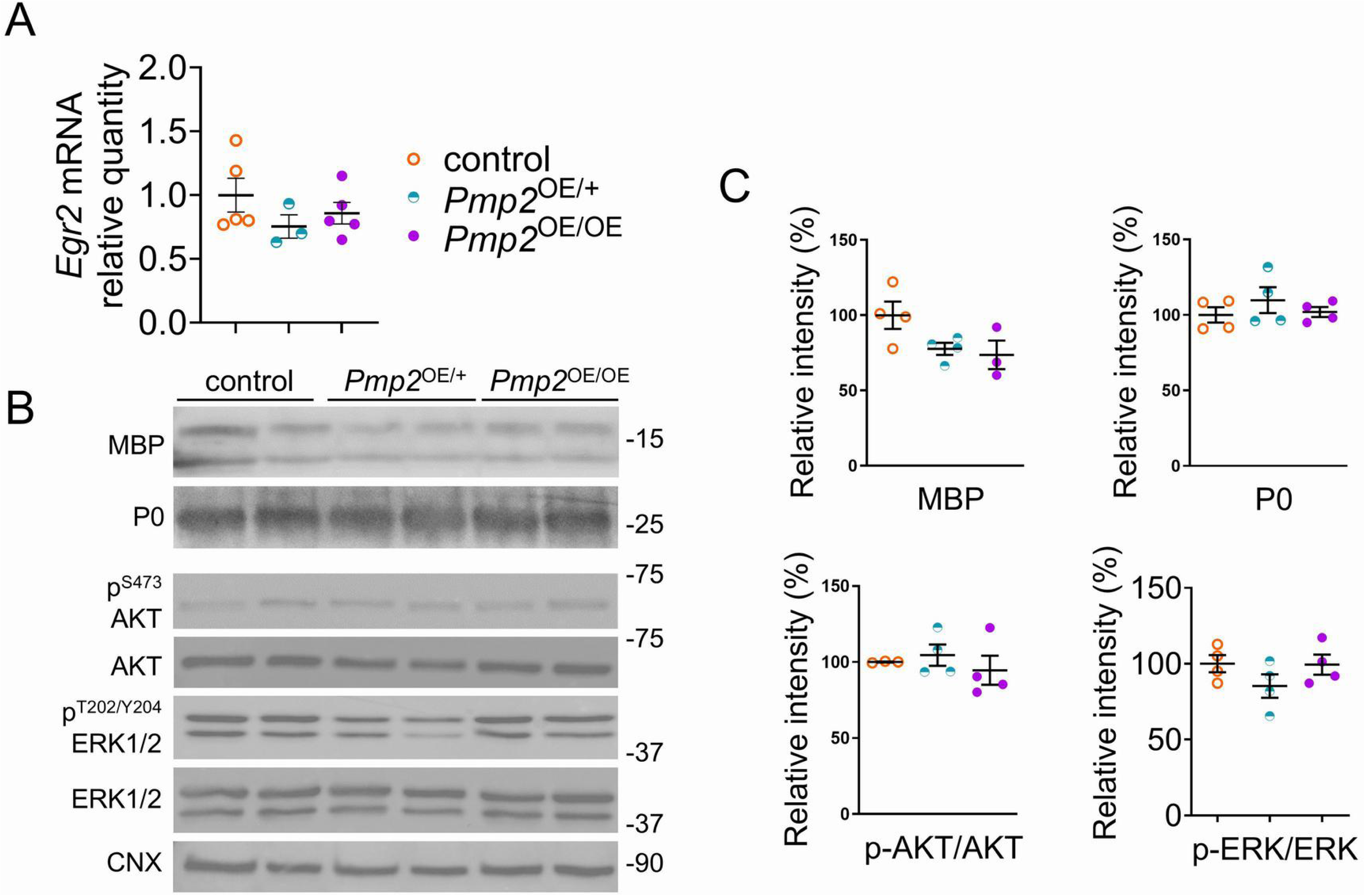
RNA and protein expression of promyelinating regulators in PMP2 overexpressing mouse sciatic nerves at P30. (A) qPCR analysis for *Egr2* mRNA level in control, *Pmp2*^OE/+^, *Pmp2*^OE/OE^ mouse sciatic nerves at P30. Error bars represent s.e.m. n = 3-5 mice, and each data point represents a different n. One-way ANOVA with Bonferroni post hoc test. (B) Western blot analysis and (C) Densitometry for MBP, P0, phosphorylated AKT (P^S473^ AKT), total AKT, phosphorylated ERK (P^T202/Y204^ ERK), and total ERK on control, *Pmp2*^OE/+^, and *Pmp2*^OE/OE^ sciatic nerves at P30 (B, C). Calnexin (CNX) was used as a protein loading control. Error bars represent s.e.m. n = 4 mice, and each data point represents a different n. One-way ANOVA with Bonferroni post hoc test.

Together, these data indicate that PMP2 overexpression increases myelin thickness without altering canonical promyelinating transcriptional programs, major myelin protein expression, or AKT/ERK signaling.

### PMP2–tdTomato is enriched in perinuclear and endoplasmic reticulum compartments without inducing ER stress

PMP2 belongs to the fatty acid-binding protein (FABP) family, members of which regulate intracellular lipid handling and membrane synthesis. However, beyond its established role as a structural myelin protein, little is known about whether PMP2 also performs a Schwann cell-specific FABP-like function in lipid trafficking. Because lipid handling and membrane synthesis are spatially compartmentalized processes ^39^, we next examined the subcellular distribution of PMP2 within the myelinating Schwann cell.

Using high-resolution imaging of teased sciatic nerve fibers, we show that PMP2 staining is enriched in the internodes in control sciatic nerve fibers, with comparable levels in the endoplasmic reticulum (ER), and only a minor fraction is localized to the nucleus (39% of internodal levels) (Fig. 5A, B). In contrast to control sciatic nerves, both *Pmp2*^OE/+^, and *Pmp2*^OE/OE^ nerves showed PMP2 enrichment in the nucleus and ER relative to internodal staining: in *Pmp2*^OE/+^, ER and nuclear PMP2 were increased by 84% and 70%, respectively, while in *Pmp2*^OE/OE^, the corresponding increases were 56% and 52% (Fig. 5A, B). The nuclear and ER enrichment of PMP2 in *Pmp2*^OE/+^, and *Pmp2*^OE/OE^ sciatic nerves persist up to 5 months of age (Fig. 5C, D).

**Figure 5.**
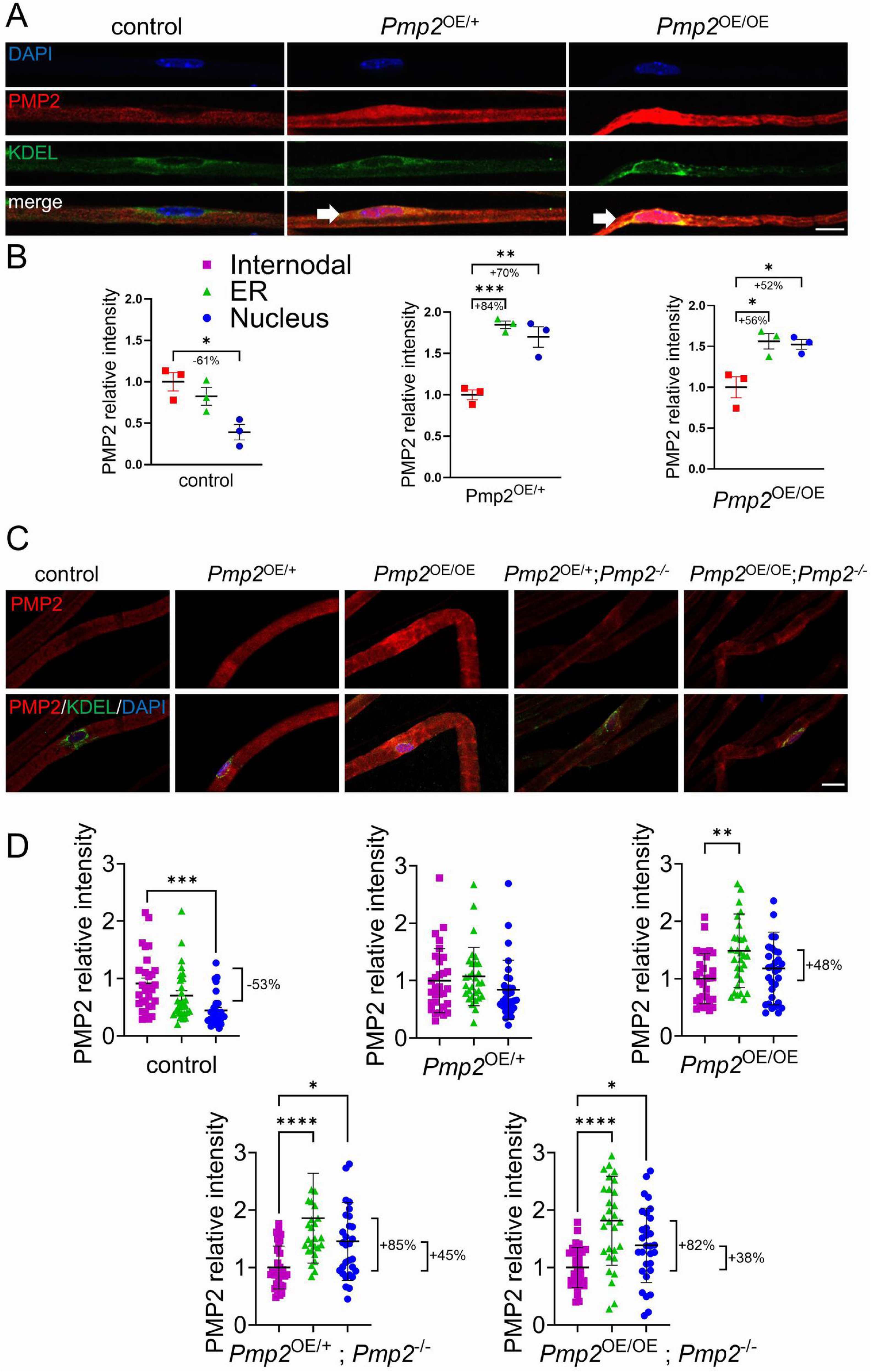
Subcellular localization of PMP2 in control and PMP2-overexpressing mouse sciatic nerves. (A) Confocal imaging of DAPI (blue), PMP2 (red), KDEL (green), in control, *Pmp2*^OE/+,^ and *Pmp2*^OE/OE^ teased fibers from mouse sciatic nerve at P30. KDEL was used as a marker of endoplasmic reticulum. Arrows indicates the colocalization of PMP2/KDEL or PMP2/DAPI. Scale bar = 10µm. (B) Relative intensity of PMP2, in the internode, nucleus, and endoplasmic reticulum in control, *Pmp2*^OE/+^, and *Pmp2*^OE/OE^ teased fibers from mouse sciatic nerve at P30. Error bars represent s.e.m. n = 3 mice, and each data point represents a different n. 10 randomized fibers per mouse were quantified. One-way ANOVA with Bonferroni post hoc test. *p<0.05, **p<0.01, ***p<0.001. (C) Confocal imaging of DAPI (blue), PMP2 (red), KDEL (green), in control, *Pmp2*^OE/+^, *Pmp2*^OE/OE^, *Pmp2*^-/-^; *Pmp2*^OE/+^, and *Pmp2*^-/-^; *Pmp2*^OE/OE^ mouse sciatic nerve teased fibers at P160. Scale bar = 10µm. (D) Relative intensity of PMP2, in the internode, nucleus, and endoplasmic reticulum in control, *Pmp2*^OE/+^, *Pmp2*^OE/OE^, *Pmp2*^-/-^; *Pmp2*^OE/+^, and *Pmp2*^-/-^; *Pmp2*^OE/OE^ mouse sciatic nerve teased fibers at P160. Error bars represent s.d. n = 31-32 fibers, and each data point represents a different n. One-way ANOVA with Bonferroni post hoc test. *p<0.05, **p<0.01, ***p<0.001, ****p<0.0001.

To determine whether this subcellular enrichment reflects is caused by increased PMP2 levels or a property specific to the PMP2-tdTomato fusion protein, we performed identical analyses in nerves expressing PMP2-tdTomato but lacking endogenous PMP2 (PMP2^OE/+^; PMP2^-/-^ and PMP2^OE/OE^; PMP2^-/-^) (Fig. 5C, D). Strikingly, PMP2-tdTomato displayed a similar nuclear and ER enrichment pattern in the absence of endogenous PMP2, indicating that this localization is likely driven by the tdTomato tag rather than by PMP2 overexpression.

Despite prominent enrichment of PMP2-tdTomato within the endoplasmic reticulum, analysis of unfolded protein response and ER stress markers revealed no induction of ER stress in *Pmp2*^OE/+^, and *Pmp2*^OE/OE^ sciatic nerves (Supplementary Fig. 3). Expression levels of canonical ER stress markers such as *Ddit3*, *Xbp-1u*, *Xbp-1s* and *Hspa5* remained unchanged across control, *Pmp2*^OE/+^, and *Pmp2*^OE/OE^ sciatic nerves, in contrast to the marked induction observed in P0^Ser63del/+^ mouse sciatic nerves, used here as a positive control sample for ER stress ^40^. These results suggest that increased PMP2-tdTomato abundance within ER-associated compartments does not perturb ER homeostasis or protein-folding capacity.

Given the nuclear and perinuclear localization of PMP2–tdTomato, we next considered whether PMP2 might influence gene expression through lipid-sensitive transcriptional pathways ^41^. However, analysis of PPAR signaling revealed no significant changes in PPAR target gene expression or pathway activation in *Pmp2*^OE/+^ and in *Pmp2*^OE/OE^ mice sciatic nerves (Supplementary Fig. 4).

In addition to perinuclear enrichment, PMP2-tdTomato was detected within non-compact myelin domains, such as Schmidt-Lanterman incisures (SLI). Our data suggests that similarly to the ER/Perinuclear enrichment, the enrichment in SLI is driven by the tdTomato tag rather than a dosage effect of increased PMP2 levels (Supplementary Fig. 5A, B).

Finally, despite the well-established efficiency of P0-Cre recombinase in myelinating Schwann cells ^26^, not all myelinating Schwann cells exhibited detectable PMP2–tdTomato signal (Supplementary Fig. 5C), suggesting that PMP2–tdTomato expression may be subject to post-transcriptional or translational regulation.

Together, these data indicate that PMP2-tdTomato localizes in the internode and non-compact myelin domains but is enriched in the ER, without inducing ER stress.

### PMP2 enhances fatty acid uptake through FABP-dependent mechanisms

The enrichment of PMP2 within metabolically active subcellular domains, including the ER, suggested that elevated PMP2 may directly influence intracellular fatty acid trafficking. To directly test this possibility, we assessed fatty acid uptake in sciatic nerves using the fluorescent long-chain fatty acid analog BODIPY ^9^.

Ex vivo analysis of teased sciatic nerve fibers revealed a significant increase in internodal BODIPY uptake in *Pmp2*^OE/+^ and in *Pmp2*^OE/OE^ mice sciatic nerves compared with teased fibers from control sciatic nerves (Fig. 6A-C). Increased fluorescence intensity was observed along the internodal regions of myelinated fibers, suggesting that the overexpression of PMP2 in Schwann cells enhances fatty acid incorporation into myelin-associated compartments *in vivo*.

**Figure 6.**
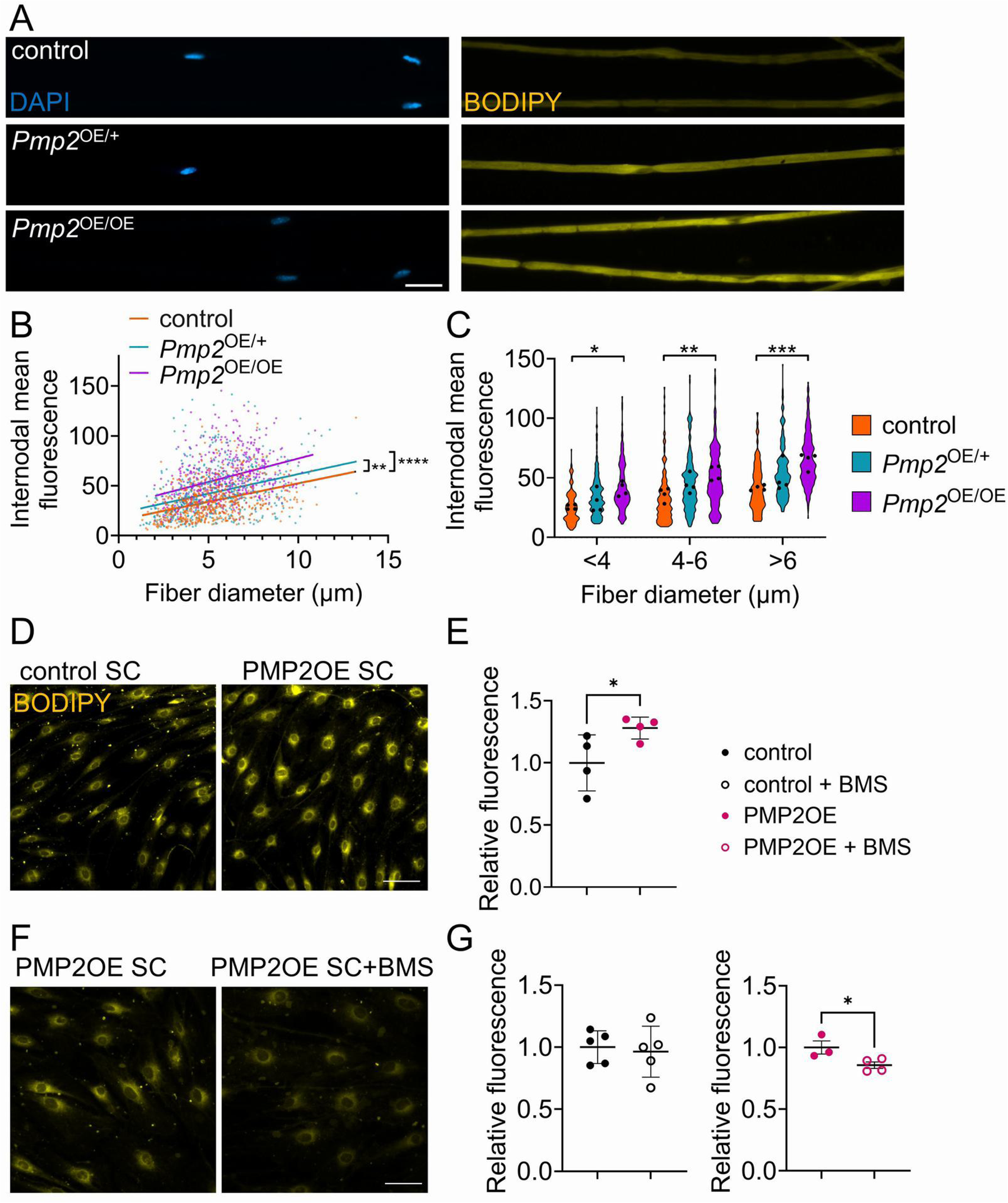
Fatty acid uptake in PMP2 overexpressing mouse sciatic nerve fibers at P15. (A) Sciatic nerve teased fibers of control, PMP2^OE/+^, and PMP2^OE/OE^ mouse sciatic nerves at P15 were treated with 5µM BODIPY-C16. Nuclei (DAPI, blue), and BODIPY (yellow) are shown. Scale bar = 20µm. (B-C) Quantification of BODIPY fluorescent signals in the internodal regions. (B) Simple linear regression. n = 500 fibers per mouse were analyzed, each data point represents a different n. (C) Violin plot of BODIPY internodal fluorescence. n = 4 mice, each data point represents a different n. Two-way ANOVA with Bonferroni post hoc. *p<0.05, **p<0.01, ***p<0.001. (D) Control and PMP2 overexpressing Schwann cells were treated with 20µM BODIPY-C12. BODIPY (yellow) is shown. Scale bar = 20µm. (E) Quantification of BODIPY fluorescent signals in Schwann cells. Error bars represent s.d. n = 4 coverslips per condition, and 100 cells were analyzed per coverslip. Each data point represents a different n. Two-tail unpaired t-test. *p<0.05. (F) PMP2 overexpressing Schwann cells were treated with 20µM BODIPY-C12 and with or without FABP inhibitor BMS309403 (BMS). BODIPY (yellow) is shown. Scale bar = 10µm. (G) Quantification of BODIPY fluorescent signals in Schwann cells. Error bars represent s.d. n = 3-5 coverslips per condition, and 100 cells were analyzed per coverslip. Each data point represents a different n. Two-tail unpaired t-test. *p<0.05.

To determine whether PMP2 overexpression in Schwann cells mediates fatty acid uptake in Schwann cells, we pharmacologically inhibited PMP2 in vitro. We used BMS309403, a small molecule that competes with fatty acids for the binding pocket of FABP family members. Using Schwann cells overexpressing PMP2 ^9^, we observe a significant increase in BODIPY uptake, consistent with the enhanced uptake observed *in vivo* (Fig. 6D, E). Importantly, treatment with BMS309403 abolished this increase, reducing fatty acid uptake to lower levels than control cells (Fig. 6F, G). These findings indicate that the enhanced fatty acid uptake associated with PMP2 overexpression is dependent on FABP-mediated transport mechanisms.

Transcriptional analysis of fatty acid transporters in *Pmp2*^OE/+^ and in *Pmp2*^OE/OE^ mice sciatic nerves revealed an upregulation of *Fatp3 Pmp2*^OE/OE^ versus control sciatic nerves (Supplementary Fig. 6), However, this finding could not be validated at the protein level due to the lack of a reliable antibody.

To determine whether PMP2 overexpression instead alters downstream mitochondrial fatty acid utilization, we next examined the expression of key regulators of mitochondrial fatty acid β-oxidation. qPCR analysis revealed no significant changes in the expression of *Cpt1a*, *Cpt2*, *Acadl*, *Acadvl*, or *Acads* in *Pmp2^OE/+^* or *Pmp2^OE/OE^* sciatic nerves compared with controls (Supplementary Fig. 7), indicating that PMP2 overexpression does not induce transcriptional remodeling of the mitochondrial β-oxidation pathway.

## Discussion

Our results demonstrate an intrinsic mechanism in which Schwann cells can promote sustained myelin thickening *in vivo*, thereby providing a new framework for peripheral myelination. Classical models have established that axon-derived cues, especially NRG1t3, are essential for initiating myelination and determining myelin thickness in the peripheral nervous system ^2,3^. However, recent studies revealed that NRG1t3 overexpression triggers lipid-centric adaptations in myelinating Schwann cells, including a major increase in PMP2 expression, and the modulation of lipid content in the myelin ^6,7,9,10^. By directly elevating PMP2 in Schwann cells, our study confirms and extends this concept by showing that enhancing Schwann cell’s lipid-handling capacity is associated with increased myelin sheath thickness *in vivo*. In other words, boosting PMP2 can phenocopy key aspects of NRG1-driven hypermyelination through a wholly glia-intrinsic route, consistent with a model in which enforced lipid availability (“force-feeding” of fatty acids) to Schwann cells promotes myelin growth.

PMP2 overexpression achieved robust hypermyelination without engaging in the classical promyelinating transcriptional program. Conventionally, NRG1t3 drives myelination through the ErbB2/3 receptor-ERK/AKT-EGR2 axis, inducing the full transcriptional program of myelin protein genes in Schwann cells ^4,5^. Any major increase in myelin thickness is generally expected to coincide with the upregulation of these pathways. However, in PMP2-overexpressing nerves, we observed decoupling of myelin growth from these canonical signals: myelin was significantly thicker even though *Egr2* mRNA and major structural myelin proteins (P0, MBP) remained at baseline levels, and neither AKT nor ERK showed sustained hyperactivation. This decoupling of morphological myelination from the canonical transcriptional response aligns with prior observations ^6,7,42,43^. For instance, exogenous NRG1t3 can drive remyelination and increase myelin thickness through PMP2 upregulation without further induction of P0, MBP, and induction of promyelinating pathways ERK and AKT ^9^. By selectively overexpressing PMP2 in Schwann cells, we induced hypermyelination, although to a lesser extent than that observed in NRG1t3-overexpressing contexts, without the need to enhance axonal NRG1 signaling or upregulate the canonical myelin transcriptional program. Thus, our results demonstrate for the first time that Schwann cells possess an intrinsic capacity *in vivo* to augment myelin growth if provided additional lipid-trafficking support, effectively breaking the long-assumed coupling between extrinsic axonal signals and achievable myelin thickness. This finding reframes the control of myelination as not purely an axon-determined process but also one shaped by Schwann cell-intrinsic and metabolic programs that can be directly modulated to enhance myelin growth.

Endogenously, PMP2 is enriched in a subset of myelinating Schwann cells, particularly those ensheathing large motor axons ^13,14^. Notably, PMP2 expression is highest during early postnatal myelination and decreases in adulthood, consistent with a role in supporting periods of active membrane growth rather than myelin maintenance. We propose that Schwann cells associated with large axons may already express high levels of PMP2 and thus near-maximal myelin thickness, whereas small-fiber Schwann cells (which are typically PMP2-negative) retain a greater dynamic range to benefit from increased lipid-trafficking capacity. This interpretation supports a threshold model for metabolic support, in which PMP2 becomes limiting only when myelin growth exceeds the normative range. In line with that idea, *Pmp2*-deficient mice exhibit only transient and mild hypomyelination, with largely preserved nerve architecture after early development ^9,14,18^, indicating that PMP2 is dispensable for baseline myelination. However, under conditions of increased myelin production, such as NRG1t3-driven hypermyelination or PMP2 overexpression, PMP2-dependent lipid-handling function becomes rate-limiting ^6,9^. This interplay raises new questions: for instance, what mechanisms normally restrict PMP2 expression to only a subset of Schwann cells ^13,14^? Are there compensatory lipid regulatory pathways in PMP2-negative Schwann cells, or do axons calibrate myelinating capacity by inducing PMP2 only in selected glia? Future studies should investigate how PMP2 is regulated *in vivo* and whether co-activation of axonal signals and glial lipid metabolism can synergistically promote remyelination beyond what either stimulus achieves alone.

We also note some unexpected enrichment for PMP2-tdTomato fusion protein to Schwann cell nuclei, the ER, or the SLI. Our data suggests those enrichments are most likely caused by the tdTomato fluorescent tag rather than a consequence of PMP2 overexpression itself. Those results are consistent with recent evidence that membrane-targeted tdTomato in Schwann cells predominantly localizes to non-compact myelin compartments (inner/outer tongues, paranodal loops, and Schmidt-Lanterman incisures) rather than compact myelin ^44^. It is possible that PMP2, as a member of the FABP family, benefits from being in the cytosolic portion of Schwann cells and therefore contributes more effectively to lipid trafficking and myelin production. Future work should determine whether sustained expression of untagged PMP2 correlates with PMP2-tdTomato localization and effect on myelination.

More broadly, our study extends the view that Schwann cells’ metabolism regulates myelin thickness. Myelination requires the synthesis and trafficking of enormous quantities of lipids, and Schwann cells must balance this biosynthetic load with energy production and nutrient uptake ^21,22^. Prior work has shown that perturbing Schwann cell lipid metabolism, through genetic disruption of lipid synthesis or β-oxidation enzymes, can limit myelin growth or lead to axon degeneration even if myelin structure initially appears intact ^45–47^. Conversely, enhancing Schwann cell metabolic output can bolster nerve function: for example, increasing glial glycolytic flux helps sustain axons after injury (*via* lactate shuttling) ^48,49^. Our findings add to this body of evidence by showing that amplifying a single Schwann cell lipid-trafficking protein can yield a durable increase in myelin thickness without external stimuli. This underscores a concept of “*adaptive myelination*”^37^ in which myelinating glia modulate myelin sheath growth in response to internal metabolic capacity and external cues.

Therapeutically, targeting glial metabolic pathways could open novel strategies for disorders of myelination. Members of the fatty-acid–binding protein family (like PMP2) have been explored as drug targets in metabolic diseases ^41^, raising the possibility that boosting a Schwann cell’s lipid-handling capability might enhance myelin repair or regeneration in neuropathies. By revealing that PMP2 acts as a dosage-sensitive facilitator linking lipid trafficking to myelin membrane expansion, our study expands the framework of myelin regulation and identifies a potential regulator for safely enhancing myelin thickness *in vivo* in Schwann cells. However it is also important to note that in humans, PMP2 may contribute to oligodendrocyte and/or astrocyte biology ^50,51^.

In summary, these findings identify PMP2 as a dosage-sensitive facilitator of myelin membrane expansion, expand the conceptual framework of myelin thickness regulation, and highlight modulation of Schwann cell lipid metabolism as a promising strategy to enhance myelin repair and regeneration alongside conventional approaches targeting axonal signaling.

## Author contributions

JH Investigation, Data analysis

CR Investigation, Data analysis

SA Investigation, Data analysis

EM Investigation, Data analysis

SE Investigation, Data analysis

KP Investigation, Data analysis

OH Investigation, Data analysis

MD Investigation

TH Investigation

BB Investigation

KM Investigation

SR Investigation

YP Conceptualization, Methodology, Investigation, Supervision, Funding, Writing-original draft

SB Conceptualization, Methodology, Investigation, Supervision, Funding, Writing-original draft

## References

1. Birchmeier, C., and Nave, K.A. (2008). Neuregulin-1, a key axonal signal that drives Schwann cell growth and differentiation. Glia 56, 1491–1497. 10.1002/glia.20753.

2. Michailov, G.V., Sereda, M.W., Brinkmann, B.G., Fischer, T.M., Haug, B., Birchmeier, C., Role, L., Lai, C., Schwab, M.H., and Nave, K.A. (2004). Axonal neuregulin-1 regulates myelin sheath thickness. Science 304, 700–703. 10.1126/science.1095862.

3. Taveggia, C., Zanazzi, G., Petrylak, A., Yano, H., Rosenbluth, J., Einheber, S., Xu, X., Esper, R.M., Loeb, J.A., Shrager, P., et al. (2005). Neuregulin-1 type III determines the ensheathment fate of axons. Neuron 47, 681–694. 10.1016/j.neuron.2005.08.017.

4. Mei, L., and Nave, K.A. (2014). Neuregulin-ERBB signaling in the nervous system and neuropsychiatric diseases. Neuron 83, 27–49. 10.1016/j.neuron.2014.06.007.

5. Newbern, J., and Birchmeier, C. (2010). Nrg1/ErbB signaling networks in Schwann cell development and myelination. Semin Cell Dev Biol 21, 922–928. 10.1016/j.semcdb.2010.08.008.

6. Belin, S., Ornaghi, F., Shackleford, G., Wang, J., Scapin, C., Lopez-Anido, C., Silvestri, N., Robertson, N., Williamson, C., Ishii, A., et al. (2019). Neuregulin 1 type III improves peripheral nerve myelination in a mouse model of congenital hypomyelinating neuropathy. Hum Mol Genet 28, 1260–1273. 10.1093/hmg/ddy420.

7. Scapin, C., Ferri, C., Pettinato, E., Zambroni, D., Bianchi, F., Del Carro, U., Belin, S., Caruso, D., Mitro, N., Pellegatta, M., et al. (2019). Enhanced axonal neuregulin-1 type-III signaling ameliorates neurophysiology and hypomyelination in a Charcot-Marie-Tooth type 1B mouse model. Hum Mol Genet 28, 992–1006. 10.1093/hmg/ddy411.

8. Fledrich, R., Stassart, R.M., Klink, A., Rasch, L.M., Prukop, T., Haag, L., Czesnik, D., Kungl, T., Abdelaal, T.A., Keric, N., et al. (2014). Soluble neuregulin-1 modulates disease pathogenesis in rodent models of Charcot-Marie-Tooth disease 1A. Nat Med 20, 1055–1061. 10.1038/nm.3664.

9. Hong, J., Garfolo, R., Kabre, S., Humml, C., Velanac, V., Roue, C., Beck, B., Jeanette, H., Haslam, S., Bach, M., et al. (2024). PMP2 regulates myelin thickening and ATP production during remyelination. Glia 72, 885–898. 10.1002/glia.24508.

10. Kim, M., Wende, H., Walcher, J., Kuhnemund, J., Cheret, C., Kempa, S., McShane, E., Selbach, M., Lewin, G.R., and Birchmeier, C. (2018). Maf links Neuregulin1 signaling to cholesterol synthesis in myelinating Schwann cells. Genes Dev 32, 645–657. 10.1101/gad.310490.117.

11. Della-Flora Nunes, G., Hong, J., Garfolo, R., Jenica, A., Mondschein, A.S., Harris, O.M., Panchal, K., Jourd’heuil, F.L., Jourd’heuil, D., Poitelon, Y., and Belin, S. (2025). PMP2 Enhances Schwann Cell Metabolism and Promotes Myelination. J Neurochem 1cS, e70265. 10.1111/jnc.70265.

12. Kozlowski, M.M., Strickland, A., Benitez, A.M., Schmidt, R.E., Bloom, A.J., Milbrandt, J., and DiAntonio, A. (2025). Pmp2+ Schwann Cells Maintain the Survival of Large-Caliber Motor Axons. J Neurosci 45. 10.1523/JNEUROSCI.1362-24.2025.

13. Yim, A.K.Y., Wang, P.L., Bermingham, J.R., Jr., Hackett, A., Strickland, A., Miller, T.M., Ly, C., Mitra, R.D., and Milbrandt, J. (2022). Disentangling glial diversity in peripheral nerves at single-nuclei resolution. Nat Neurosci 25, 238–251. 10.1038/s41593-021-01005-1.

14. Zenker, J., Stettner, M., Ruskamo, S., Domenech-Estevez, E., Baloui, H., Medard, J.J., Verheijen, M.H., Brouwers, J.F., Kursula, P., Kieseier, B.C., and Chrast, R. (2014). A role of peripheral myelin protein 2 in lipid homeostasis of myelinating Schwann cells. Glia 62, 1502–1512. 10.1002/glia.22696.

15. Abe, M., Makino, A., Murate, M., Hullin-Matsuda, F., Yanagawa, M., Sako, Y., and Kobayashi, T. (2021). PMP2/FABP8 induces PI(4,5)P(2)-dependent transbilayer reorganization of sphingomyelin in the plasma membrane. Cell Rep 37, 109935. 10.1016/j.celrep.2021.109935.

16. Ruskamo, S., Yadav, R.P., Sharma, S., Lehtimaki, M., Laulumaa, S., Aggarwal, S., Simons, M., Burck, J., Ulrich, A.S., Juffer, A.H., et al. (2014). Atomic resolution view into the structure-function relationships of the human myelin peripheral membrane protein P2. Acta Crystallogr D Biol Crystallogr 70, 165–176. 10.1107/S1399004713027910.

17. Uusitalo, M., Klenow, M.B., Laulumaa, S., Blakeley, M.P., Simonsen, A.C., Ruskamo, S., and Kursula, P. (2021). Human myelin protein P2: from crystallography to time-lapse membrane imaging and neuropathy-associated variants. FEBS J 288, 6716–6735. 10.1111/febs.16079.

18. Stettner, M., Zenker, J., Klingler, F., Szepanowski, F., Hartung, H.P., Mausberg, A.K., Kleinschnitz, C., Chrast, R., and Kieseier, B.C. (2018). The Role of Peripheral Myelin Protein 2 in Remyelination. Cell Mol Neurobiol 38, 487–496. 10.1007/s10571-017-0494-0.

19. Hong, Y.B., Joo, J., Hyun, Y.S., Kwak, G., Choi, Y.R., Yeo, H.K., Jwa, D.H., Kim, E.J., Mo, W.M., Nam, S.H., et al. (2016). A Mutation in PMP2 Causes Dominant Demyelinating Charcot-Marie-Tooth Neuropathy. PLoS Genet 12, e1005829. 10.1371/journal.pgen.1005829.

20. Motley, W.W., Palaima, P., Yum, S.W., Gonzalez, M.A., Tao, F., Wanschitz, J.V., Strickland, A.V., Loscher, W.N., De Vriendt, E., Koppi, S., et al. (2016). De novo PMP2 mutations in families with type 1 Charcot-Marie-Tooth disease. Brain 139, 1649–1656. 10.1093/brain/aww055.

21. Montani, L., Pereira, J.A., Norrmen, C., Pohl, H.B.F., Tinelli, E., Trotzmuller, M., Figlia, G., Dimas, P., von Niederhausern, B., Schwager, R., et al. (2018). De novo fatty acid synthesis by Schwann cells is essential for peripheral nervous system myelination. J Cell Biol 217, 1353–1368. 10.1083/jcb.201706010.

22. Poitelon, Y., Kopec, A.M., and Belin, S. (2020). Myelin Fat Facts: An Overview of Lipids and Fatty Acid Metabolism. Cells 9. 10.3390/cells9040812.

23. Boucanova, F., and Chrast, R. (2020). Metabolic Interaction Between Schwann Cells and Axons Under Physiological and Disease Conditions. Front Cell Neurosci 14, 148. 10.3389/fncel.2020.00148.

24. Beirowski, B., Babetto, E., Golden, J.P., Chen, Y.J., Yang, K., Gross, R.W., Patti, G.J., and Milbrandt, J. (2014). Metabolic regulator LKB1 is crucial for Schwann cell-mediated axon maintenance. Nat Neurosci 17, 1351–1361. 10.1038/nn.3809.

25. Nave, K.A. (2010). Myelination and support of axonal integrity by glia. Nature 4c 8, 244–252. 10.1038/nature09614.

26. Feltri, M.L., D’Antonio, M., Previtali, S., Fasolini, M., Messing, A., and Wrabetz, L. (1999). P0-Cre transgenic mice for inactivation of adhesion molecules in Schwann cells. Ann N Y Acad Sci 883, 116–123.

27. Poitelon, Y., Kozlov, S., Devaux, J., Vallat, J.M., Jamon, M., Roubertoux, P., Rabarimeriarijaona, S., Baudot, C., Hamadouche, T., Stewart, C.L., et al. (2012). Behavioral and molecular exploration of the AR-CMT2A mouse model Lmna (R298C/R298C). Neuromolecular Med 14, 40–52. 10.1007/s12017-012-8168-z.

28. Mondschein, A.S., DiPersio, M.R., Zajaceskowski, J., Nimmagadda, H., Acheta, J., Salinero, A.E., Haslam, S., Poitelon, E., Elston, S., McFarland, E., et al. (2025). High-Fat Diet Disrupt Nerve Function by Targeting Schwann Cells. J Peripher Nerv Syst 30, e70036. 10.1111/jns.70036.

29. Jeanette, H., Marziali, L.N., Bhatia, U., Hellman, A., Herron, J., Kopec, A.M., Feltri, M.L., Poitelon, Y., and Belin, S. (2021). YAP and TAZ regulate Schwann cell proliferation and differentiation during peripheral nerve regeneration. Glia cS, 1061-1074. 10.1002/glia.23949.

30. Moore, S.M., Jeong, E., Zahid, M., Gawron, J., Arora, S., Belin, S., Sim, F., Poitelon, Y., and Feltri, M.L. (2024). Loss of YAP in Schwann cells improves HNPP pathophysiology. Glia 72, 1974–1984. 10.1002/glia.24592.

31. Hong, J., Kirkland, J.M., Acheta, J., Marziali, L.N., Beck, B., Jeanette, H., Bhatia, U., Davis, G., Herron, J., Roue, C., et al. (2024). YAP and TAZ regulate remyelination in the central nervous system. Glia 72, 156–166. 10.1002/glia.24467.

32. Poitelon, Y., Matafora, V., Silvestri, N., Zambroni, D., McGarry, C., Serghany, N., Rush, T., Vizzuso, D., Court, F.A., Bachi, A., et al. (2018). A dual role for Integrin alpha6beta4 in modulating hereditary neuropathy with liability to pressure palsies. J Neurochem 145, 245–257. 10.1111/jnc.14295.

33. Catignas, K.K., Frick, L.R., Pellegatta, M., Hurley, E., Kolb, Z., Addabbo, K., McCarty, J.H., Hynes, R.O., van der Flier, A., Poitelon, Y., et al. (2021). alpha(V) integrins in Schwann cells promote attachment to axons, but are dispensable in vivo. Glia cS, 91-108. 10.1002/glia.23886.

34. Weaver, M.R., Shkoruta, D., Pellegatta, M., Berti, C., Palmisano, M., Ferguson, S., Hurley, E., French, J., Patel, S., Belin, S., et al. (2025). The STRIPAK complex is required for radial sorting and laminin receptor expression in Schwann cells. Cell Rep 44, 115401. 10.1016/j.celrep.2025.115401.

35. Acheta, J., Hong, J., Jeanette, H., Brar, S., Yalamanchili, A., Feltri, M.L., Manzini, M.C., Belin, S., and Poitelon, Y. (2022). Cc2d1b Contributes to the Regulation of Developmental Myelination in the Central Nervous System. Front Mol Neurosci 15, 881571. 10.3389/fnmol.2022.881571.

36. Follis, R., Prabhu, V.V., and Carter, B.D. (2025). The Influence of Schwann Cell Metabolism and Dysfunction on Axon Maintenance. Glia 73, 2338–2352. 10.1002/glia.70071.

37. Knowles, J.K., Batra, A., Xu, H., and Monje, M. (2022). Adaptive and maladaptive myelination in health and disease. Nat Rev Neurol 18, 735–746. 10.1038/s41582-022-00737-3.

38. Salzer, J., Feltri, M.L., and Jacob, C. (2024). Schwann Cell Development and Myelination. Cold Spring Harb Perspect Biol 16. 10.1101/cshperspect.a041360.

39. Barnes-Velez, J.A., Aksoy Yasar, F.B., and Hu, J. (2023). Myelin lipid metabolism and its role in myelination and myelin maintenance. Innovation (Camb) 4, 100360. 10.1016/j.xinn.2022.100360.

40. D’Antonio, M., Musner, N., Scapin, C., Ungaro, D., Del Carro, U., Ron, D., Feltri, M.L., and Wrabetz, L. (2013). Resetting translational homeostasis restores myelination in Charcot-Marie-Tooth disease type 1B mice. J Exp Med 210, 821–838. 10.1084/jem.20122005.

41. Furuhashi, M., and Hotamisligil, G.S. (2008). Fatty acid-binding proteins: role in metabolic diseases and potential as drug targets. Nat Rev Drug Discov 7, 489–503. 10.1038/nrd2589.

42. Domenech-Estevez, E., Baloui, H., Meng, X., Zhang, Y., Deinhardt, K., Dupree, J.L., Einheber, S., Chrast, R., and Salzer, J.L. (2016). Akt Regulates Axon Wrapping and Myelin Sheath Thickness in the PNS. J Neurosci 36, 4506–4521. 10.1523/JNEUROSCI.3521-15.2016.

43. Sheean, M.E., McShane, E., Cheret, C., Walcher, J., Muller, T., Wulf-Goldenberg, A., Hoelper, S., Garratt, A.N., Kruger, M., Rajewsky, K., et al. (2014). Activation of MAPK overrides the termination of myelin growth and replaces Nrg1/ErbB3 signals during Schwann cell development and myelination. Genes Dev 28, 290–303. 10.1101/gad.230045.113.

44. Reinert, A., Winkler, U., Goebbels, S., Komarek, L., Möbius, W., Zanker, H.S., Fledrich, R., Stassart, R.M., Hirrlinger, P.G., Nave, K.-A., et al. (2026). Red fluorescent labeling of myelin by membrane-targeted tdTomato in transgenic mouse lines. bioRxiv, 2026.2004.2017.718425. 10.64898/2026.04.17.718425.

45. Norrmen, C., Figlia, G., Lebrun-Julien, F., Pereira, J.A., Trotzmuller, M., Kofeler, H.C., Rantanen, V., Wessig, C., van Deijk, A.L., Smit, A.B., et al. (2014). mTORC1 controls PNS myelination along the mTORC1-RXRgamma-SREBP-lipid biosynthesis axis in Schwann cells. Cell Rep 9, 646-660. 10.1016/j.celrep.2014.09.001.

46. Viader, A., Sasaki, Y., Kim, S., Strickland, A., Workman, C.S., Yang, K., Gross, R.W., and Milbrandt, J. (2013). Aberrant Schwann cell lipid metabolism linked to mitochondrial deficits leads to axon degeneration and neuropathy. Neuron 77, 886–898. 10.1016/j.neuron.2013.01.012.

47. Chung, H.L., Wangler, M.F., Marcogliese, P.C., Jo, J., Ravenscroft, T.A., Zuo, Z., Duraine, L., Sadeghzadeh, S., Li-Kroeger, D., Schmidt, R.E., et al. (2020). Loss- or Gain-of-Function Mutations in ACOX1 Cause Axonal Loss via Different Mechanisms. Neuron 106, 589–606 e586. 10.1016/j.neuron.2020.02.021.

48. Babetto, E., Wong, K.M., and Beirowski, B. (2020). A glycolytic shift in Schwann cells supports injured axons. Nat Neurosci 23, 1215–1228. 10.1038/s41593-020-0689-4.

49. Jha, M.K., Lee, Y., Russell, K.A., Yang, F., Dastgheyb, R.M., Deme, P., Ament, X.H., Chen, W., Liu, Y., Guan, Y., et al. (2020). Monocarboxylate transporter 1 in Schwann cells contributes to maintenance of sensory nerve myelination during aging. Glia 68, 161–177. 10.1002/glia.23710.

50. Gargareta, V.-I., Mougios, N., Hahn, E.T., Siems, S.B., Crisp, S.J., Varga, B., Karadottir, R.T., Buescher, J.M., Jung, R.B., Ramesh, V., et al. (2026). Fatty acid binding protein-8 (FABP8/PMP2) reveals molecular heterogeneity of myelin sheaths in the human CNS. bioRxiv, 2026.2006.2009.730898. 10.64898/2026.06.09.730898.

51. Zhang, Y., Chen, K., Sloan, S.A., Bennett, M.L., Scholze, A.R., O’Keeffe, S., Phatnani, H.P., Guarnieri, P., Caneda, C., Ruderisch, N., et al. (2014). An RNA-sequencing transcriptome and splicing database of glia, neurons, and vascular cells of the cerebral cortex. J Neurosci 34, 11929–11947. 10.1523/JNEUROSCI.1860-14.2014.

