## Supplemental files for "FABP8/PMP2 is a positive regulator of PNS myelination"

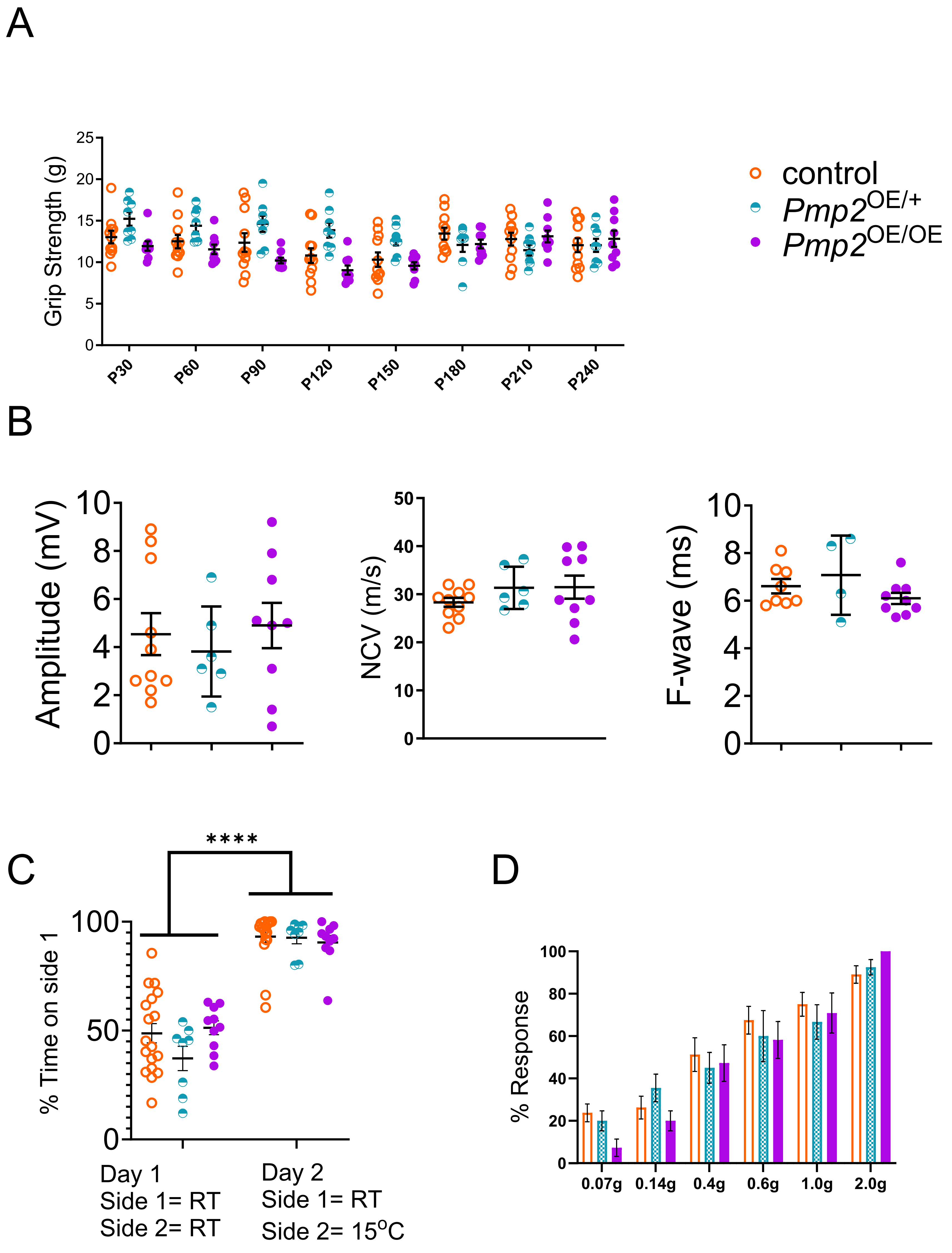


**Supplementary Figure 1. Functional assessment in PMP2 overexpressing mice.** (A) Longitudinal measurement of motor function in control, *Pmp2*$^{OE/+}$, and *Pmp2*$^{OE/OE}$ mice is assessed using the grip strength assay from P30 to P240. Males and females were used. Error bars represent s.e.m. n= 8-11 mice, and each data point represents a different n. (B) Electrophysiological analysis in control, *Pmp2*$^{OE/+}$, and *Pmp2*$^{OE/OE}$ mice at P240. Compound muscle action potential amplitude, nerve conduction velocity, and F-wave latency were measured. Error bars represent s.e.m. n = 4-10 mice, and each data point represents a different n. (C) Thermal preference of control, *Pmp2*$^{OE/+}$, and *Pmp2*$^{OE/OE}$ mice at P240 was assessed and reported as time spent on side 1 (RT) for both days. Males and females were used. Error bars represent s.e.m. n= 8-11 mice, and each data point represents a different n. (D) Mechanical threshold of control, *Pmp2*$^{OE/+}$, and *Pmp2*$^{OE/OE}$ mice at P240. Males and females were used. Error bars represent s.e.m. n= 6-18 mice. (B) One-way ANOVA with Bonferroni post hoc test. (A, C, D) Two-way ANOVA with Bonferroni post hoc test. ****p<0.0001.

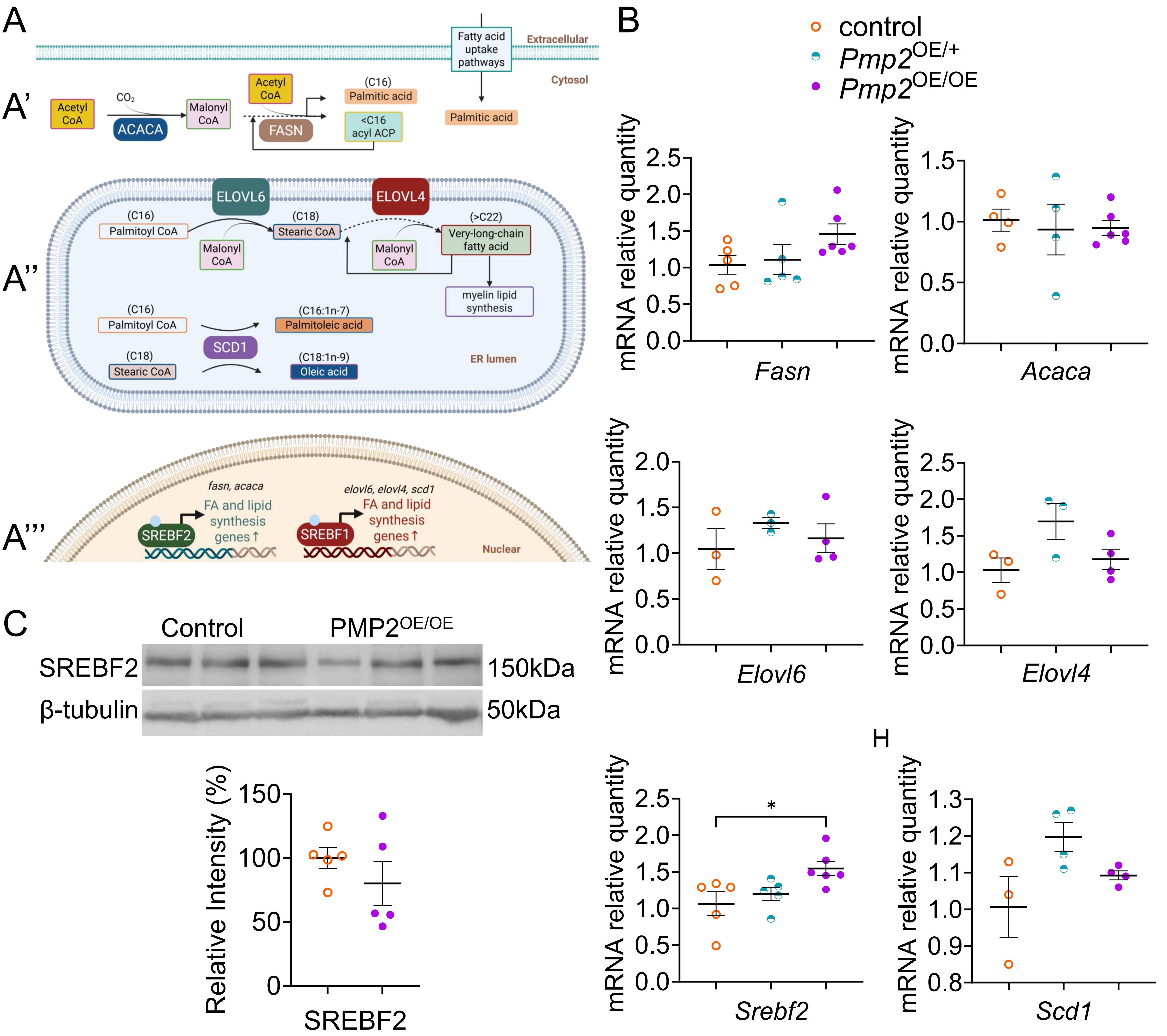


**Supplementary figure 2. Expression of fatty acid synthesis and lipogenesis regulators in PMP2 overexpressing mouse sciatic nerves at P30.** (A) Schematic of short-chain, long-chain, very-long-chain, and monounsaturated fatty acid synthesis pathways. (A') In the cytosol, ACACA converts acetyl-CoA and $CO_2$ into malonyl-CoA, and FASN elongates malonyl-CoA into palmitate (C16). Palmitate can be synthesized de novo or taken up from extracellular sources and used in the ER for further elongation. (A'') Long-chain and very-long-chain fatty acids are synthesized in the ER lumen. ELOVL6 adds malonyl-CoA to palmitoyl-CoA to generate C18 stearic-CoA, and successive cycles by ELOVL4 elongate the chain to very-long-chain fatty acids, which are incorporated into myelin lipids. (A'') Mono-unsaturated fatty acid are also synthesized in the ER lumen. SCD1 converts palmitoyl-CoA into palmitoleic acid and stearic-CoA into oleic acid. (A''') *Fasn* and *Acaca* are regulated by SREBF2, whereas *Elovl6*, *Elovl4*, and *Scd1* are regulated by SREBF1. (B) qPCR analysis for *Fasn, Acaca, Evolv6, Evolv4, Srebf2* and *Scd1* mRNA level in control, *Pmp2*$^{OE/+}$, *Pmp2*$^{OE/OE}$ mouse sciatic nerves at P30. *Rpl27* was used as housekeeping control. Error bars represent s.e.m. n = 3-6 mice, and each data point represents a different n. One-way ANOVA with Bonferroni post hoc test. *$p < 0.05$. (C) Western blot analysis and densitometry for SREBF2 in control and *Pmp2*$^{OE/OE}$ sciatic nerves at P30. β-tubulin was used as a protein loading control. Error bars represent s.e.m. n = 5 mice, and each data point represents a different n. Two-tailed unpaired t-test.

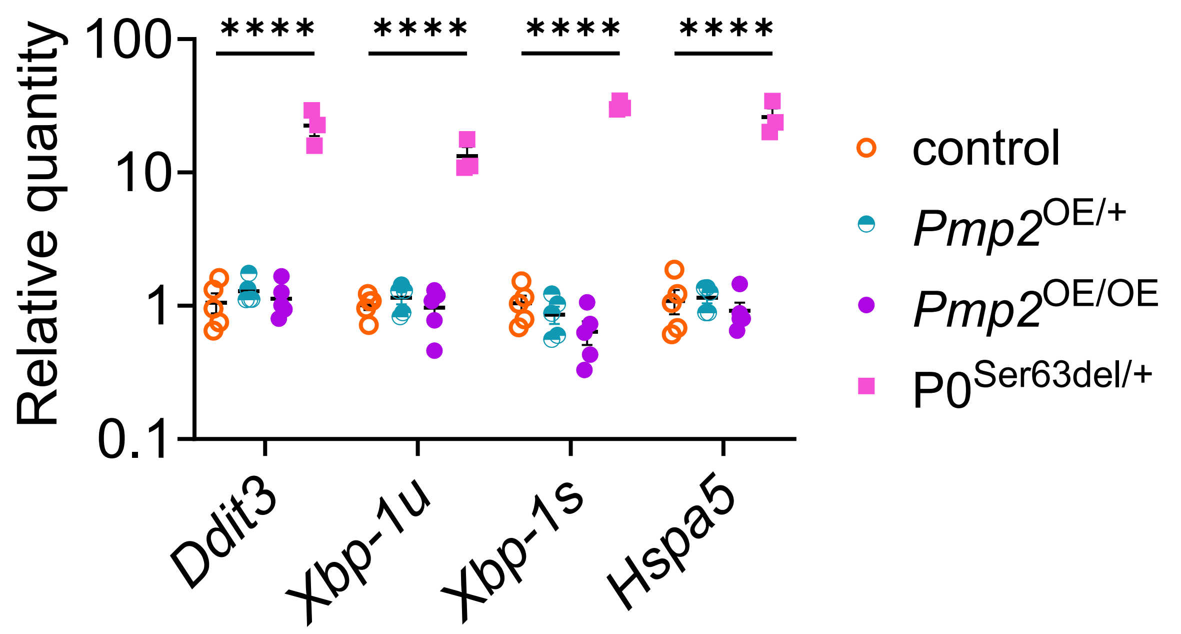


**Supplemental Figure 3. qPCR analysis of ER stress markers in PMP2 overexpressing mouse sciatic nerves at P30.** qPCR analysis for *Ddit3*, *Xbp-1u*, *Xbp-1s* and *Hspa5* mRNA level in control, *Pmp2*$^{OE/+}$, *Pmp2*$^{OE/OE}$ and P0$^{ser63del/+}$ mouse sciatic nerves at P30. 18S was used as a housekeeping control. *Ddit3*, *Xbp-1u*, *Xbp-1s* and *Hspa5* were used as a marker of ER stress. P0$^{ser63del/+}$ was used as a positive control for Schwann cell ER stress. Error bars represent s.e.m. n = 3 mice, and each data point represents a different n. Two-way ANOVA with Bonferroni post hoc test. ****p<0.0001.

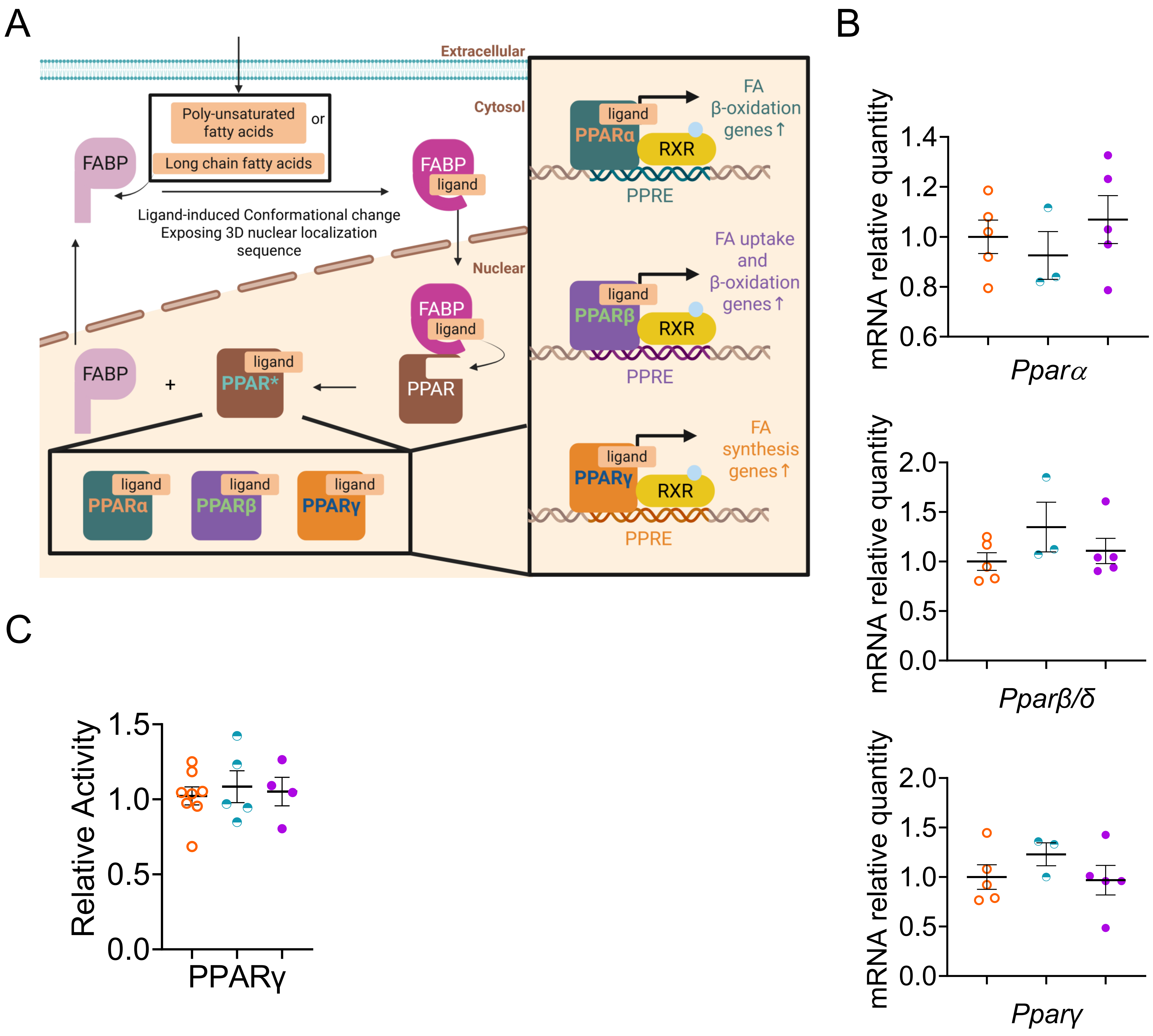


**Supplemental Figure 4. FABP-mediated activation of PPAR transcription factors in PMP2 overexpressing mouse sciatic nerves at P30.** (A) Schematic of FABP-mediated activation of PPARs. FABPs translocate to the nucleus with their ligands and deliver them to PPARs, thereby activating PPAR transcriptional activity. PPARα promotes transcription of fatty acid β-oxidation genes. PPARβ/δ promotes transcription of fatty acid uptake and β-oxidation genes. PPARγ promotes transcription of fatty acid synthesis genes. (B) qPCR analysis for *Pparβ/δ, Pparα*, and *Pparγ* mRNA level in control, *Pmp2*$^{OE/+}$, *Pmp2*$^{OE/OE}$ mouse sciatic nerves at P30. *Rpl27* was used as housekeeping control. Error bars represent s.e.m. n = 3-5 mice, and each data point represents a different n. One-way ANOVA with Bonferroni post hoc test. (C) ELISA analysis for PPARγ activity in the control, PMP2$^{OE/+}$ and PMP2$^{OE/OE}$ mouse sciatic nerves at P30. Error bars represent s.e.m. n ≥ 4 mice, and each data point represents a different n. Two-tail unpaired t-test.

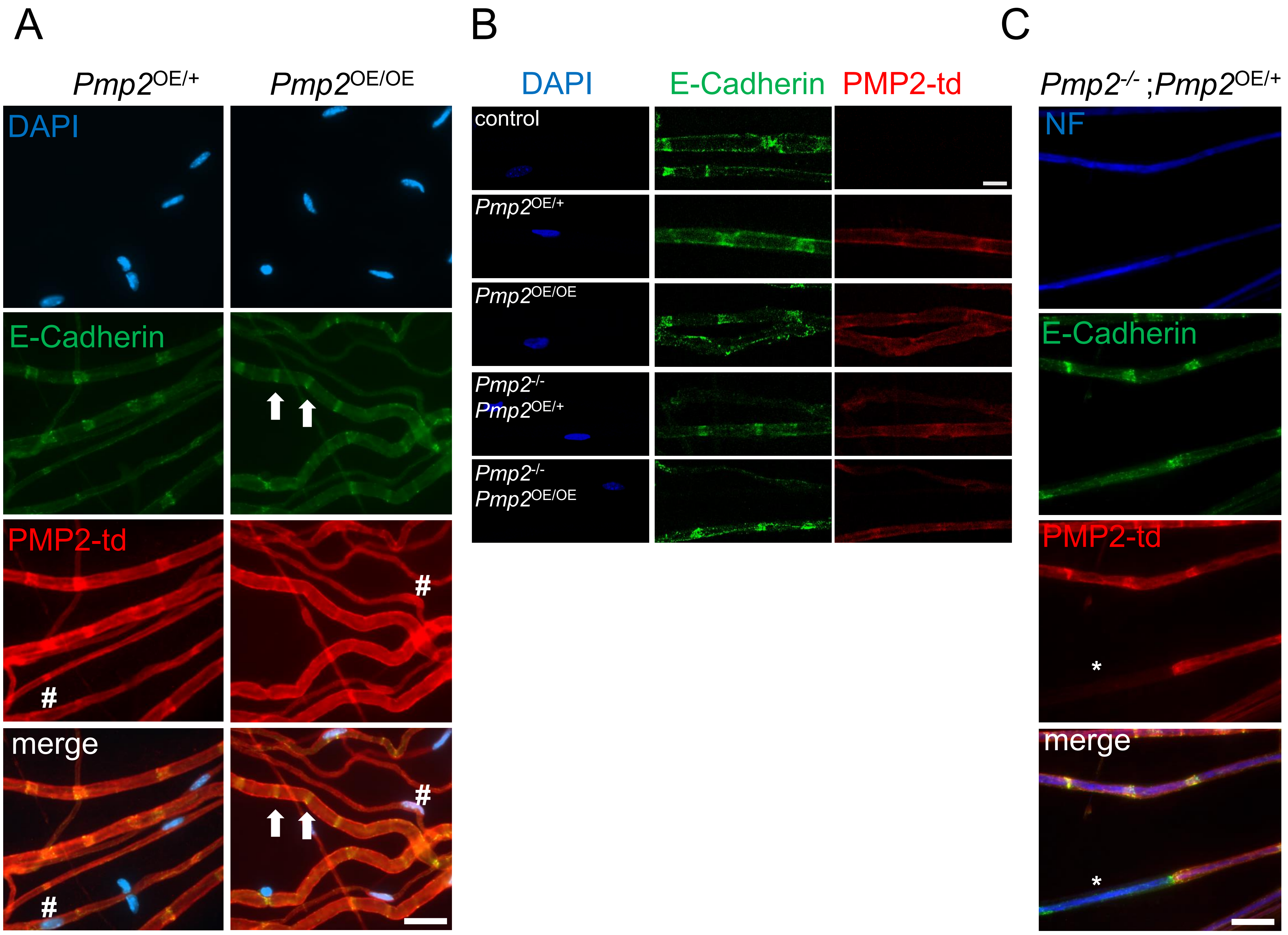


**Supplemental Figure 5. Subcellular localization of PMP2 in control and PMP2-overexpressing mouse sciatic nerves.** (A) Epifluorescent imaging of DAPI (blue), E-cadherin (green), and PMP2-tdTomato (red) in *Pmp2*$^{OE/+}$ and *Pmp2*$^{OE/OE}$ teased fibers from mouse sciatic nerve at P30. E-Cadherin was used as a marker of non-compact myelin. Arrows indicate Schmidt-Lanterman incisures (SLI, non-compact myelin regions), and hashtags indicate nuclear and perinuclear regions. Scale bars = 20μm. (B) Confocal imaging of DAPI (blue), E-cadherin (green), and PMP2 protein (red) in control, *Pmp2*$^{OE/+}$, *Pmp2*$^{OE/OE}$, *Pmp2*$^{-/-}$; *Pmp2*$^{OE/+}$, and *Pmp2*$^{-/-}$; *Pmp2*$^{OE/OE}$ mouse sciatic nerve teased fibers at P160. Scale bars = 10μm. (C) Epifluorescent imaging of Neurofilament (NF, blue), E-Cadherin (green), and PMP2-tdTomato (red), and in *Pmp2*$^{-/-}$;*Pmp2*$^{OE/+}$ teased fibers from mouse sciatic nerve at P160. NF was used as a marker for axons. The image shows two Schwann cells myelinating the same axon, one expressed PMP2-TdTomato, and one does not. Scale bar = 20μm.

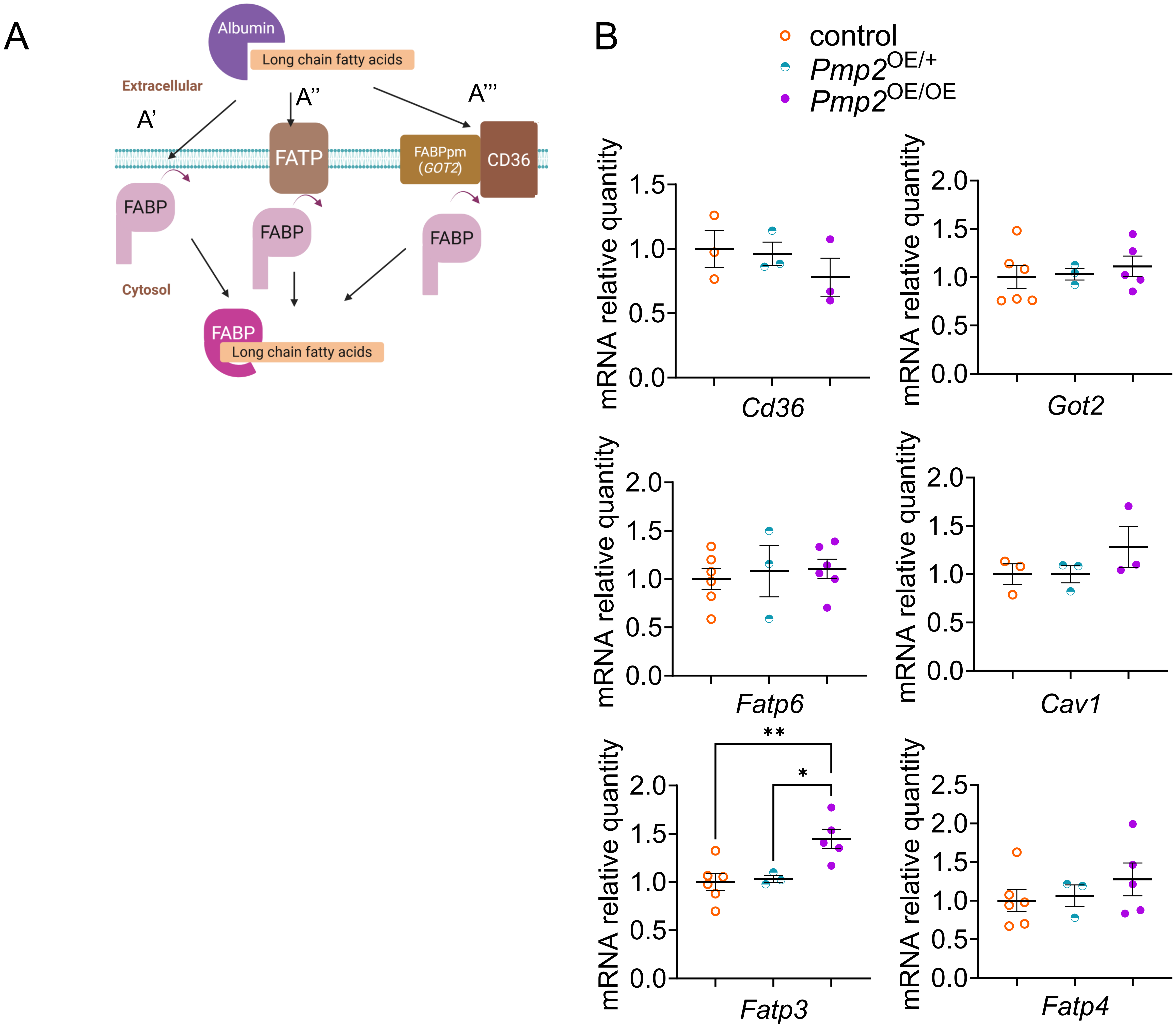


**Supplementary Figure 6. Expression of fatty acid uptake regulators in PMP2 overexpressing mouse sciatic nerves at P30.** (A) Schematic of fatty acid uptake mediated by fatty acid-binding proteins (FABPs). (A') Direct collision of FABPs with the plasma membrane allows uptake from the membrane. (A'') FABPs can also mediate fatty acid uptake through fatty acid transporters (FATPs). (A''') Scavenger receptor CD36 and membrane-bound FABPpm facilitate fatty acid uptake, and FABPs interact with CD36 via collision. (B) qPCR analysis for *CD36, Got2, Fatp3, Fatp4, Fatp6* and *Cav1* mRNA level in control, *Pmp2*$^{OE/+}$, *Pmp2*$^{OE/OE}$ mouse sciatic nerves at P30. *Rpl27* was used as housekeeping control. CAV1 is known to regulate the activity of CD36. Error bars represent s.e.m. n = 3-6 mice, and each data point represents a different n. One-way ANOVA with Bonferroni post hoc test. *p < 0.05, *p < 0.01.

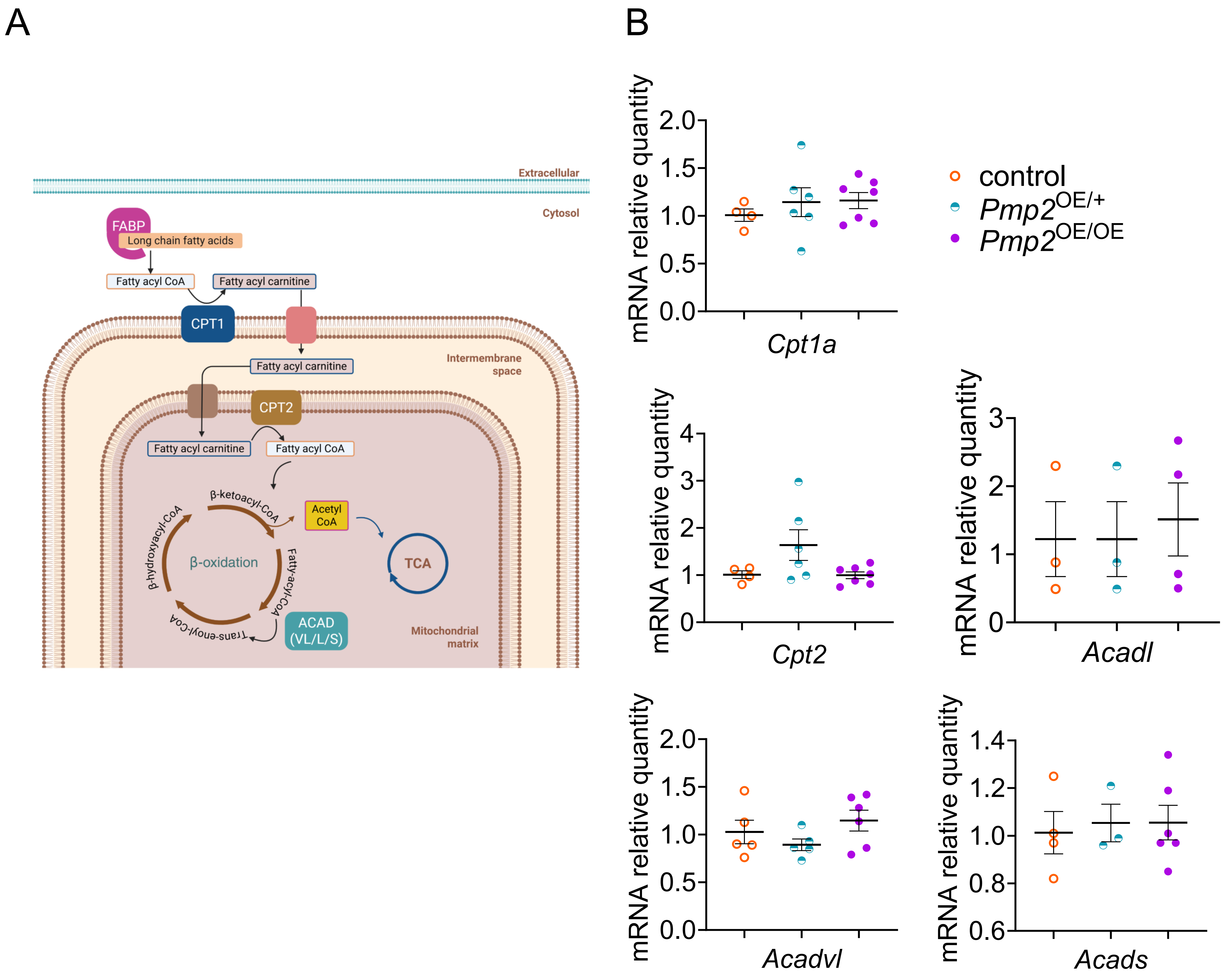


**Supplemental Figure 7. Expression of mitochondrial fatty acid β-oxidation regulators in PMP2 overexpressing mouse sciatic nerves at P30.** (A) Schematic of mitochondrial fatty acid β-oxidation. FABPs deliver long-chain fatty acids to mitochondria. The rate-limiting step of mitochondrial fatty acid β-oxidation is catalyzed by CPT1, which converts fatty-acyl-CoA into fatty-acyl carnitine. CPT2 transports fatty-acyl carnitine across the mitochondrial inner membrane into the matrix. Fatty-acyl carnitine is converted back to fatty-acyl-CoA, which enters the β-oxidation cycle. β-Oxidation is catalyzed by acyl-CoA dehydrogenases: ACADS for short-chain, ACADL for long-chain, and ACADVL for very-long-chain fatty acids. (B) qPCR analysis for *Cpt1a, Cpt2, Acadl, Acadvl* and *Acads* mRNA level in control, *Pmp2*$^{OE/+}$, *Pmp2*$^{OE/OE}$ mouse sciatic nerves at P30. *Rpl27* was used as housekeeping control. Error bars represent s.e.m. n = 3-7 mice, and each data point represents a different n. One-way ANOVA with Bonferroni post hoc test.
